# Where top-down meets bottom-up: Cell-type specific connectivity map of the whisker system

**DOI:** 10.1101/2023.07.31.551377

**Authors:** N. Rault, T. Bergmans, N. Delfstra, BJ Kleijnen, F. Zeldenrust, T. Celikel

## Abstract

Sensorimotor computation integrates bottom-up world state information with top-down knowledge and task goals to form action plans. In the rodent whisker system, a prime model of active sensing, evidence shows neuromodulatory neurotransmitters shape whisker control, affecting whisking frequency and amplitude. Since neuromodulatory neurotransmitters are mostly released from subcortical nuclei and have long-range projections that reach the rest of the central nervous system, mapping the circuits of top-down neuromodulatory control of sensorimotor nuclei will help to systematically address the mechanisms of active sensing. Therefore, we developed a neuroinformatic target discovery pipeline to mine the Allen Institute’s Mouse Brain Connectivity Atlas. Using network connectivity analysis, we identified new putative connections along the whisker system and anatomically confirmed the existence of 42 previously unknown monosynaptic connections. Using this data, we updated the sensorimotor connectivity map of the mouse whisker system and developed the first cell-type-specific map of the network. The map includes 157 projections across 18 principal nuclei of the whisker system and neuro-modulatory neurotransmitter-releasing. Performing a graph network analysis of this connectome, we identified cell-type specific hubs, sources, and sinks, provided anatomical evidence for monosynaptic inhibitory projections into all stages of the ascending pathway, and showed that neuromodulatory projections improve network-wide connectivity. These results argue that beyond the modulatory chemical contributions to information processing and transfer in the whisker system, the circuit connectivity features of the neuromodulatory networks position them as nodes of sensory and motor integration.

## 1 Introduction

In the wild, whisking rodents are subterranean mammals that live in dark and narrow environments. They heavily rely on the rhythmic protraction and retraction of their whiskers to navigate their natural habitat. Organized as closed-loop circuits (**Kleinfeld et al., 1999**), distributed networks in the brain control whisking as an adaptive sensorimotor computation while they encode sensory information originating from the sensory periphery and compute the future position of the whiskers as an iterative process (**Voigt et al., 2008**, **Voigt et al., 2015**, **McElvain et al., 2018**).

The neural circuits that contribute to the sensory representations acquired through whisker contacts and the positional controls of whiskers were last summarized more than a decade ago **(Bosman et al., 2011)**. Building on *(1)* the first comprehensive sensorimotor circuit map in the whisker system (**Kleinfeld et al., 1999**), *(2)* the fundamental experimental work that described the lemniscal and paralemniscal ascending neural circuits (**Chial et al., 1991**, **Lu and Lin, 1992**), and *(3)* the descending projections originating from the motor cortex (**Ashwell, 1982**, **Hemelt and Keller, 2008**), Bosman and colleagues described the connectome between 43 nuclei contributing to the sensorimotor integration in the whisker system. In particular, 12 of the nuclei described in their studies as well as 6 brainstem regions that release neuro-modulatory transmitters showed 74 connections (**Supplemental Figure 1**, for the complete network **Supplemental Figure 2**).

Over the last decade, new experimental studies have provided direct evidence that the network connectivity along the whisker system is more expansive than previously shown. For example, a study by **Jeong et al., 2016**, focusing on the motor nuclei in the mouse, in particular MOp (M1), revealed projections to PO and VPM, as well as to the contralateral PSV, SPV, and VTA. (**Munõz-Castañeda et al., 2021**) confirmed both this last connection and the monosynaptic Mop (M1) projections to PPN and SPVi. In parallel, direct monosynaptic projections originating from bfd and targeting TM, RT (nRT), and ZI (**Zakiewicz et al., 2014**), ZI projections to VII (**Takatoh et al., 2021**), VII innervation of SPVi (**Bellavance et al., 2017**), VPM to Mop (M1) connections (**Hooks et al., 2013**) and bidirectional communication between Mop (M1) and RT (nRT) were shown (**Yang et al., 2022**).

New monosynaptic projections along the sensorimotor pathways were also discovered to originate from several neuromodulatory neurotransmitter releasing nuclei. **Hosp et al., 2011** identified a projection between the VTA and Mop (M1), showing the contribution of dopaminergic connections originating from the VTA in motor skill learning. Other studies showed that the VTA serves as a hub of divergence with projections to SI, LC, DR, RT (nRT) (**Dunigan et al., 2021**), and bfd (**Aransay et al., 2015**), all critical for the information processing in the whisker system. Finally, the novel discoveries along the neuromodulatory circuits and their integration with nuclei in the whisker system are not limited to the dopaminergic system. Monosynaptic cholinergic projections to RT (nRT) (**Sokhadze et al., 2018**), SI projections to ZI (**Zhou et al., 2018**), ZI projections to VTA (**An et al., (2021)**), reciprocal connections between PO and SC (**Benavidez et al., 2021**), SI projections to PSV, VTA, VPM, SI (**Benavidez et al., 2021**) and DR (**Benavidez et al., 2021**, **Dorocic et al., (2014)**) all provide anatomical evidence for neuromodulatory contributions to sensorimotor control.

The newly found connections mentioned above require an update of the previously drawn connectivity map of the rodent whisker system. In this study, by leveraging the connectivity data in the Allen Mouse Brain Atlas (**Oh et al., 2014**), we update the previously reported connectome (**Bosman et al., 2011**), develop the first-generation cell-type specific connectivity map along the mouse whisker system and investigate the contribution of neuromodulator-releasing nuclei to top-down regulation and information propagation in this network. In addition to the nuclei from the ascending somatosensory pathway and the motor cortex, we consider the following networks in our updated whisker system connectivity map: The facial motor nucleus (VII), as it contains the motoneurons innervating the muscle situated in the whisker pad (**Ashwell, 1982**). The Superior Colliculus (SC), as it is targeted by connections from the trigeminal nuclei and targets VII (**Hemelt and Keller, 2008**). The reticular nucleus of the thalamus (RT (nRT)) is added as the main source of inhibition of the thalamic nuclei (**Pinault et al., 1995**). The Zona Incerta (ZI) is included in the map for its role in the paralemniscal pathway (**Urbain and Deschênes, 2007a**) and its somatotopic map representation of whiskers (**Nicolelis et al., 1992**).

By analyzing 732 anterograde tracing experiments performed in 95 wild-type and transgenic lines using the open-source toolbox we developed (https://github.com/DepartmentofNeurophysiology/NeuralNet-ABC), we determine the connectivity along the ascending, descending, and neuro-modulator pathways. We show that the sensorimotor circuits in the whisker system include 157 projections; 42 of these projections had not been described in the literature previously, but are confirmed anatomically herein. Additionally, 40 connections are identified as plausible monosynaptic projections and reported here as putative connections. By performing graph network analysis, we further identify cell-type specific hubs, sources, and sinks in this connectome and, finally, show that neuromodulatory projections improve network-wide connectivity.

## 2 Materials & Methods

### 2.1 Experimental pipeline

All experimental data (https://connectivity.brain-map.org/) and our custom-written toolbox for analysis (https://github.com/DepartmentofNeurophysiology/NeuralNet-ABC) are available online. Experimental procedures for data acquisition have been detailed by the Allen Brain Institute before (**Oh et al., 2014**). In short, genetically encoded fluorescent proteins were expressed in anatomically targeted regions using cell-type-specific or non-specific promoters in transgenic and wild-type mice. The resulting anterograde projection labeling was visualized using serial two-photon tomography. The data were projected onto a 3D normal brain built upon a volumetric average template from 1,675 wild-type mice with 10 *µm* isotropic voxel resolution. The 3D reference brain was then parcellated into 43 isocortical areas, 329 subcortical gray matters structures, 81 fiber tracts, and 8 ventricular structures for each hemisphere to identify the main anatomical origin and target areas based on a standard atlas (**Wang et al., 2020**). For each structure, four variables were extracted: the volume, the intensity, the density, and the energy. The volume represents the volume of tracer solution in mm^3^ present in a brain structure; the intensity is the sum of the brightness of every pixel in a given structure (the tracer producing a fluorescent reaction); the density is the number of pixels considered as expressing the gene divided by the total number of pixels present in the structure, and the energy is the sum of pixel intensities divided by the number of pixels in the structure. The results were stored in a JSON format containing a list of structures with their associated extracted variables. To build our connectivity matrices, we first downloaded the data using Allen Brain Institute’s API (**Figure 1**), and identified experiments that targeted 57 target nuclei, which have been experimentally identified as a part of the sensorimotor circuits in the rodent brain (**Supplemental Figure 1**). From these experiments, we filtered out those with less than 50% of viral solution injected in the targeted area (i.e., where more than half of the solution missed the targeted area). Each experiment was saved in a JSON file containing a list of structures (723 per hemisphere) and the measures of the projection volume, intensity, density, and energy. Next, we grouped the experiments per transgenic line and projected the data from each experiment onto a 723×1446 matrix (723 structures for each hemisphere), with the sources of the experiments as rows and the structures receiving the connections as columns (**Figure 1**).

**Fig. 1.**
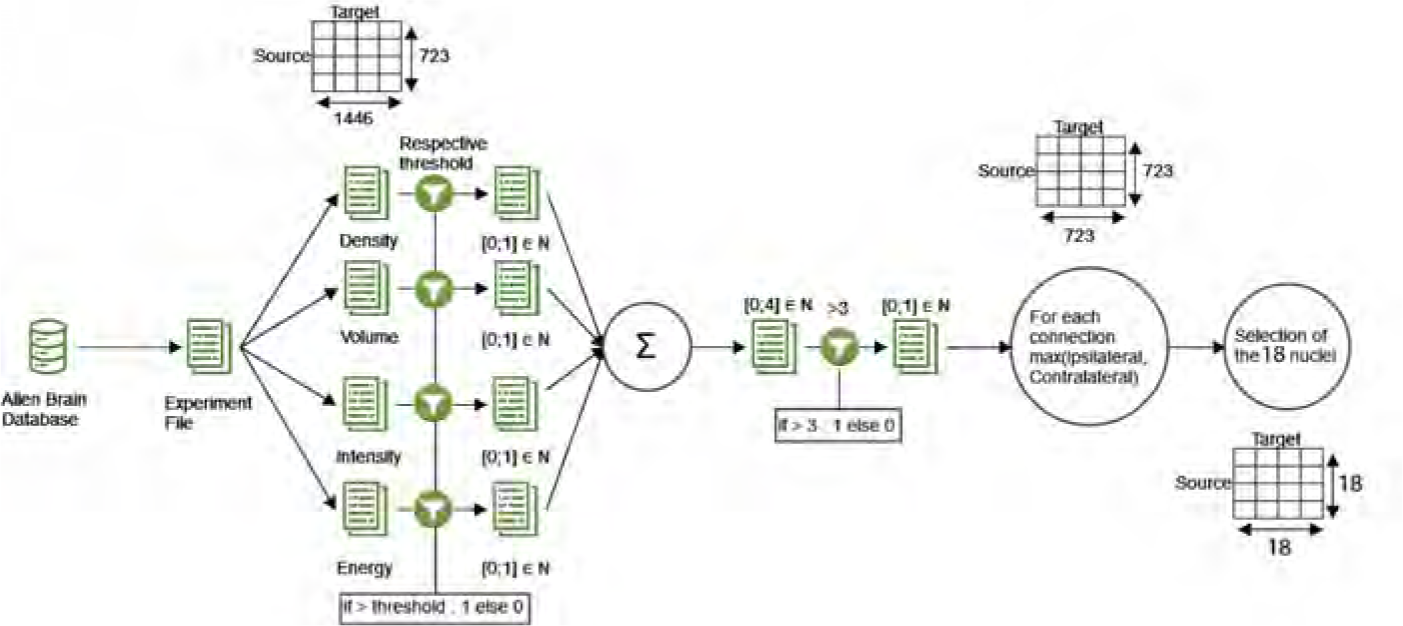
The data processing pipeline. For each transgenic line, the data were downloaded from the Allen Brain database. Each experiment file was split into 4 matrices of size 723×1446 (see section 2.2), one per variable (density, volume, intensity, and energy). A threshold specific to each variable (see Connectivity Validation) was applied and 4 matrices with values of either zero or one were retrieved. These matrices were summed and a threshold equal to 4 was applied, so that only the connections that satisfied the conditions on all 4 variables remained. This way, a matrix composed of zeros and ones was obtained. Due to the sparse nature of contralateral hemispheric projections, which will increase the false negative rate, the connectomes across the hemispheres were merged by selecting the highest value. From the resulting matrix of size 723×723, we retrieved the connections for the nuclei of interest (see Supplemental Figure 1.b and 1.c). This resulted in a binary connectivity matrix with 324 (18×18) possible edges.

### 2.2 Specificity Calculation

To quantify the percentage of solution injected in the target area, we downloaded for each experiment a 3D representation of the animal’s brain, in which every voxel’s value represents the density of solution within the area with the voxel’s coordinate. The area of injection was manually annotated by the Allen Brain Institute, and every voxel’s value outside of the injection zone was set to 0. We downloaded a 3D mask of the injection structure for each experiment. This consists of a 3D matrix representing the mouse brain where only the voxel within the coordinates of the targeted structure is equal to one, and every voxel outside of the structure is set to 0. We performed a pairwise multiplication between the mask and the injection density matrix, giving us only the part of the injection within the targeted structure. We binarized the result by setting every voxel above 0 to 1 and did the same for the injection density. We finally divided the masked injection density (binarized injection density that has been multiplied by the mask) by the binarized injection density. These results give us the percentage of injection within the targeted area, called specificity. As stated in the experimental pipeline, we used this value to filter out experiments with less than 50% specificity. To validate the newly found connections, we relied on the Allen Brain Institute specificity calculation and accepted only the anatomical validation from experiments with a specificity above 75%. This value is obtained through the same process as described previously, only without binarization of the voxel’s value.

### 2.3 Transgenic categorization

Transgenic lines were considered “excitatory” if presented as such in the literature or if the marked cells were glutamatergic, i.e. for example expressed glutamate or vesicular glutamate transporters and other proteins necessary for glutamatergic signaling. “Inhibitory” cells were classified as cells expressing Gamma-Aminobutyric Acid (GABA), glycine neurotransmitters, or other proteins related to inhibitory signaling, such as GABA receptors. Transgenic lines were classified as “uncategorized” if marked cells expressed both excitatory and inhibitory proteins, or if they expressed neither (e.g., dopamine, adrenaline, serotonin, or other neuromodulatory proteins with no direct excitatory or inhibitory effect). See **Supplemental Table 1** for the list and the corresponding citations.

### 2.4 Connectivity validation

To validate the connections identified in the connectivity matrices derived from the Allen Brain Institute’s experiments, we compared them to a subset (N=34) of connections previously reported by Bosman et al. (2011) between a subset of 18 nuclei discussed in the introduction. For each variable, we determined a variable-specific threshold by systematically increasing the threshold until one of the 34 reference connections was no longer detected in the wild-type experiments. The last value for which we found 100% of true positives was considered the correct threshold. This process yielded four distinct thresholds, which were applied to each transgenic line (density threshold: 9 × 10*^−^*^5^ (a.u.), intensity threshold: 222 (a.u.), energy threshold: 2891 × 10*^−^*^2^ (a.u.), volume threshold: 395 × 10*^−^*^4^ (in mm^3^)).

A connection was considered valid only if it appeared on each variable-specific matrix after threshold application. This approach ensured that we retained previously known connections while avoiding assumptions about the underlying connectivity that might have arisen from minimizing false positives. By focusing solely on true positives, we did not impose any preconceived notions about the pattern of connectivity between different nuclei. Cross-validating our connections with the four available variables minimized the likelihood of false negatives. As an additional safeguard, we verified that the genes marked by the transgenic lines in the Allen Brain Institute’s experiments were indeed expressed in the targeted structure using the transgenic characterization tool available on the Allen Brain Institute’s website. Applying these thresholds resulted in four connectivity matrices: one for the wild type and one for each transgenic characterization (excitatory, inhibitory, or uncategorized), combined in one (**Figure 2.b and Supplemental Figure 3**) and compiled into connectivity maps (**Figure 2.a and 3, Supplemental Figure 4 and 5**). To corroborate the connections, calculations were repeated across animal lines. We reviewed the literature and identified new connections from our results (**Figure 4**). The density of these connections was normalized for every connection source so that the total density projected from a node sums to 1. We then analyzed the distribution of outgoing density amongst targeted nodes to assess the sparsity of the projections originating from each nucleus (**Figure 5 and 6**). New connections were validated through anatomical confirmation using the imaging data available in the Allen Brain Database from experiments with at least 75% of the viral solution injected in the targeted structure as calculated by the Allen Brain Institute. Finally, we constructed a summary map that includes previously established and newly discovered connections (**Figure 4 and 7**). For anatomical validation of the identified connections, we superimposed the results from the 2-photon imaging experiments onto a mean model of the mouse brain. The two images for each connection were manually aligned using fiducial markers such as the hippocampus or the outline of the brain. The limitations of this approach are discussed in the discussion section. To summarize, in order to identify new connections, we 1) used our calculated connectivity map as a “hypothesis generation” tool to find likely projections across the connectivity map using experimental data with 50% specificity, which means that at least half of the injection volume is inside the target nucleus boundary. We 2) identified all experiments that meet a secondary criterion (i.e., 75% or better specificity), anatomically confirmed the data, and provided the data as supplemental material (**Supplemental Figures 6 to 35**). And finally, 3) we built a summary matrix based on the results of Step 2 with “anatomically confirmed” connections. The remaining projections identified in Step 1 are presented as “putative connections”.

**Fig. 2.**
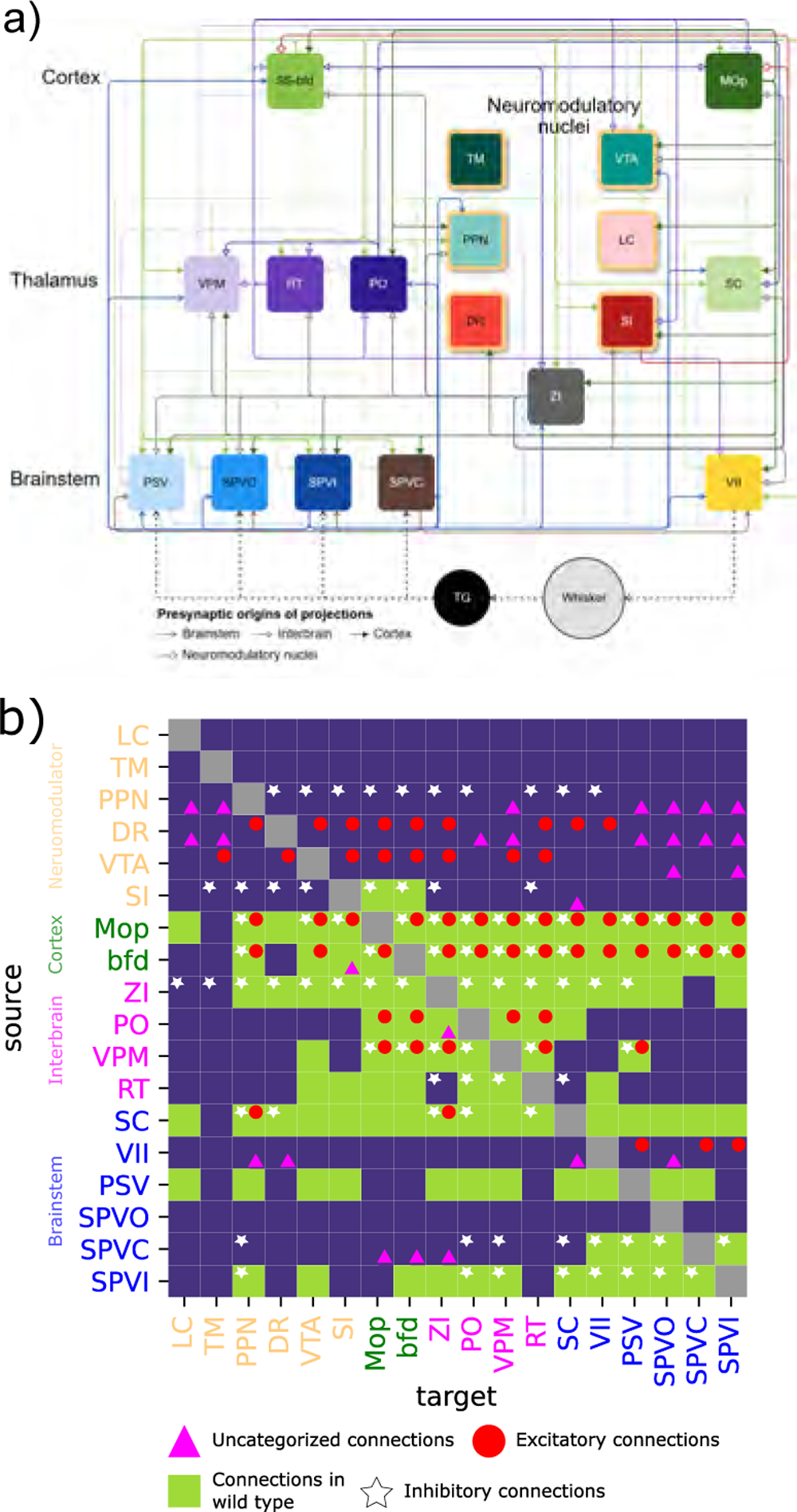
Sensorimotor connectivity map of the whisker system in wild-type mice. **a)** Connectivity map across 18 nuclei with 108 connections computed from wild-type experiments in the Allen Brain database. A dashed line represents a flow of information outside of the central nervous system. The neuromodulatory structures are represented with yellow borders. **b**) Binary representation of the connectivity matrix combining wild, excitatory, inhibitory and unclassified transgenic lines. LC, TM, PPN, and DR were not targeted in wild-type animals, thus their projections to the rest of the network are not shown. See Figure 7 for the updated connectome.

### 2.5 Network analysis

Based on the obtained connectivity maps, we analyzed the network properties of neuromodulator-releasing nuclei and their contribution to the overall connectivity of the sensorimotor circuit. To assess the small-world nature of our network, we compared our summary graph to an Erdos-Renyi (ER) graph. First, we determined the total number of connections in our summary map and calculated the probability of a connection. We then constructed an ER graph with the same number of edges, nodes, and connection probability. Using the

GRETNA toolbox (**Wang et al., 2015**), we computed the clustering coefficient and the shortest path for both the original and the ER graph. According to Watts and Strogatz (**1998**), a small-world network exhibits higher clustering values and a lower shortest path length compared to ordered and random networks. We therefore considered our summary network to be a small-world network if its clustering coefficient was higher than that of the ER graph and its average shortest path was lower than that of the ER graph.

In addition to the ER network, we constructed a summary graph without the neuromodulator-releasing nuclei. We then computed the nodal clustering coefficient, the nodal shortest path, the degree centrality, the nodal efficiency, the nodal local efficiency, and the betweenness centrality for both networks. To determine whether the neuromodulator-releasing nuclei had an impact on the network structure, we compared the distributions of these values using a permutation test with 10,000 permutations.

Finally, we assessed the presence of hubs in the extended network (57×57, **Supplemental Figure 2**). To do that, we first computed the out-degree as the sum of output and the in-degree as the sum of input for each node. The sum of those two variables gives the degree centrality. We considered every node with a degree centrality above the average plus one standard deviation as a hub (**Figure 6**).

## 3 Results

We analyzed data from 162 experiments on wild-type mice and 570 experiments on 94 different transgenic lines. Of these 732 experiments, 501 presented at least 50% specificity. Focusing on the 18-nuclei subnetwork, out of the 18 × 18 = 324 possible connections for each type of mouse, 237 connections were identified in the wild-type strain. Experiments on the transgenic lines revealed 151 excitatory connections, 198 inhibitory connections, and 211 uncategorized connections exhibiting values above 0 without threshold application. Among these 797 connections, 44% (i.e., 351 projections) met the criteria for anatomical connections (see Materials and Methods for details) and originated from experiments with at least 50% specificity. For the 18-nuclei sub-network, **Bosman et al., 2011** observed 74 connections (**Supplemental Figure 1.b, 1.c**). Our study identified 108 connections in the wild type (**Figure 2.a, 2.b**) and 243 connections in transgenic lines. The identified connections are distributed across the different cell-type-specific connectivity maps. Among these connections, 58 are excitatory (**Figure 2.b; Supplemental Figure 4**), 84 are inhibitory (**Figure 2.b; Figure 3**), and 101 are uncategorized (**Figure 2; Supplemental Figure 5**), with 27 of these not appearing on either excitatory or inhibitory connectivity maps (**Figure 2**).

**Fig. 3.**
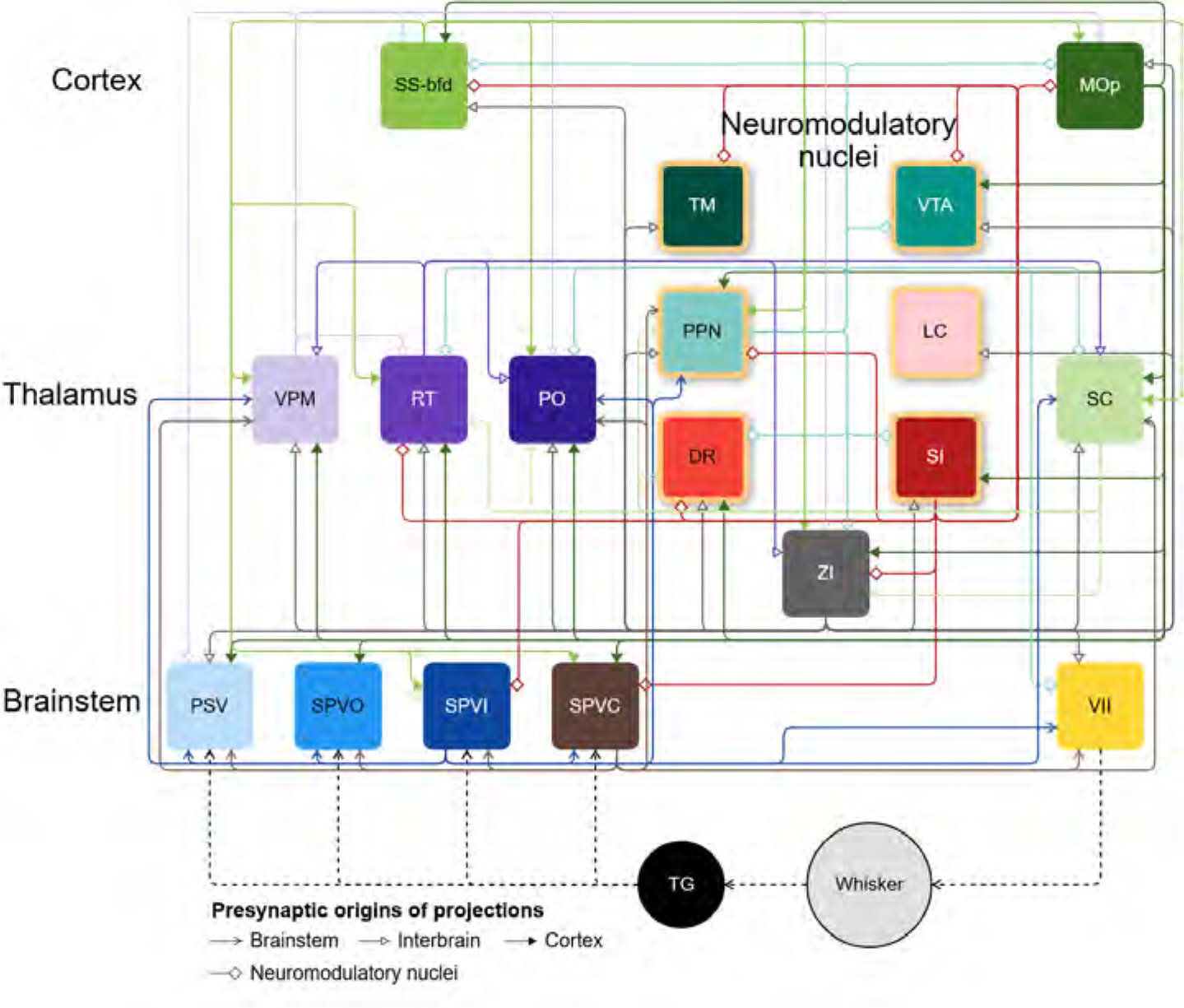
The connectome of the inhibitory projections in the mouse whisker system. The connectivity map includes 84 connections. A dashed line represents the flow of information outside of the central nervous system. The nodes with yellow borders are neuromodulatory structures.

Among the structures presented in this study, the tuberomammillary nucleus, the locus coeruleus, and the oral part of the spinal nucleus have not been targeted by experiments in the Allen Brain database, resulting in a lack of connectivity data for these structures. Additionally, on the wild-type map, connectivity data is not available for the tuberomammillary nucleus, the pedunculopontine nucleus, the locus coeruleus, and the dorsal nucleus raphe.

In the next sections, we will discuss the connections originating from each of the 18 nuclei, validated by our methods.

**Fig. 4.**
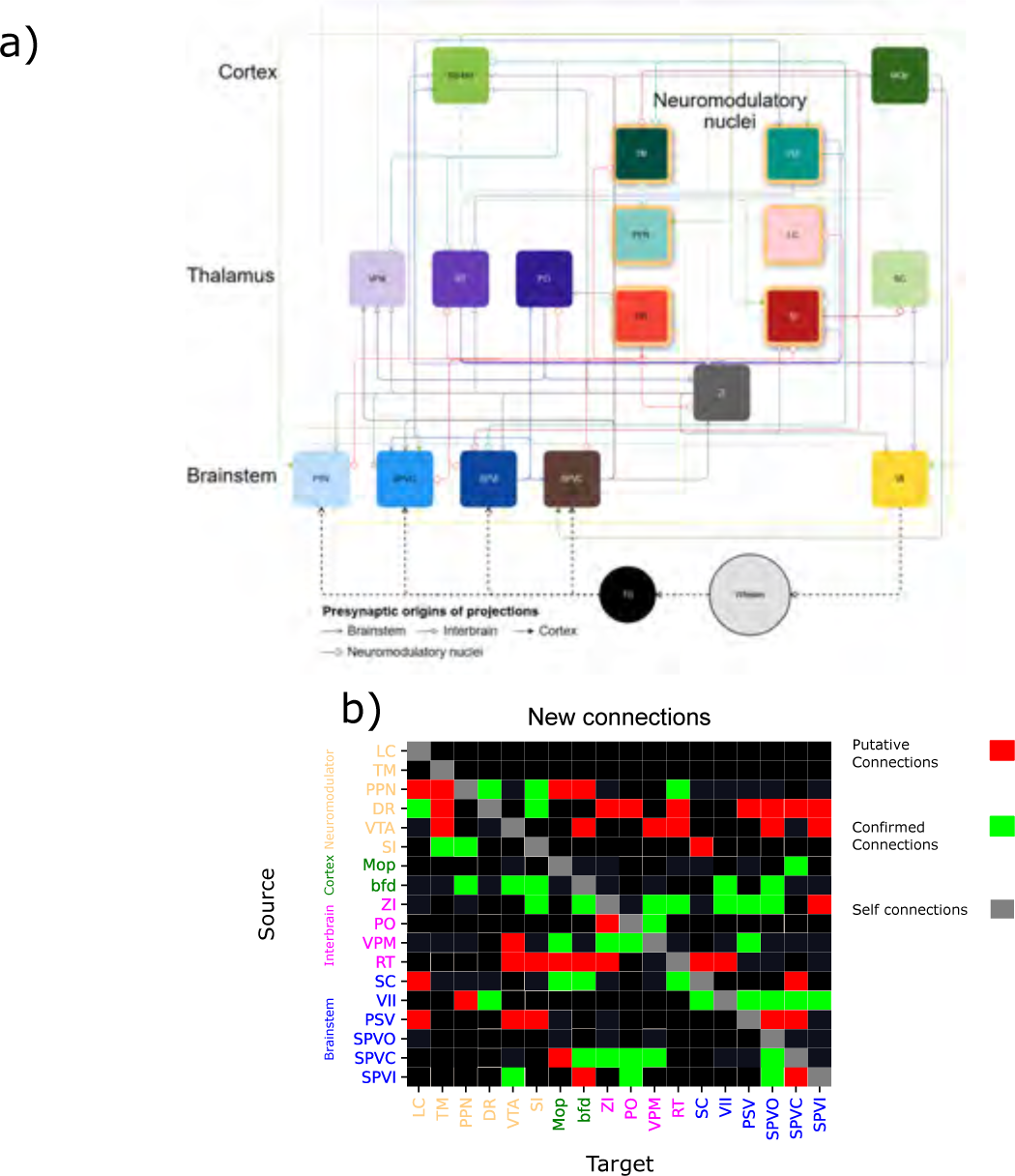
Newly discovered monosynaptic projections across the sensorimotor map of the mouse whisker system. **a)** A graphical representation of the connectome. A dashed line represents the flow of information outside of the central nervous system. The neuromodulatory structures are represented with yellow borders. **b)** 42 of the 82 identified in the experiments from the Allen Brain database were anatomically confirmed and included in the summary map (Figure 7). The vertical axis represents the source of the connection, and the horizontal axis the target. Red cells indicates putative connections, green cells anatomically validated connections, black cells represent previously reported connections or lack of connections whereas gray represent self-connections that were not considered in this study.

### 3.1 Trigeminal nuclei

The information originating from whisker contact with a surface propagates along the brainstem, the thalamus, and the primary somatosensory cortex while making closed-loop connections with the motor system (**Kleinfeld et al., 1999**). This information is not solely transmitted via excitatory connections, as inhibition is observed early in the sensory pathway (**Kleinfeld et al., 1999**). In line with this, our findings reveal the presence of inhibitory presynaptic cells in SPVi and SPVc. This suggests that both inhibitory and excitatory activity play a role in processing tactile information as early as the brainstem (**Figure 2**).

Our findings reveal an inhibitory connection originating from SPVi targeting SC. The inhibitory nature of synaptic projections from SPVi to trigeminal nuclei and VII suggests an inhibitory control of touch processing already at the brainstem level. Moreover, SPVi inhibitory projections to PO and VPM, further emphasize the brainstem’s role in whisker information processing. The presence of this ascending inhibition implies that SPVi not only relays information but also filters it. Additionally, SPVi inhibits PPN, hinting at a potential role in modulating cholinergic neurons. Connections toward VTA, bfd, ZI were identified in wild-type mice.

Connections to VTA, (PO) and SPVo were anatomically confirmed, the connections toward bfd and SPVc remain putative.

SPVi is not the only structure that targets cholinergic neurons. We demonstrated that inhibitory presynaptic cells in the trigeminal complex target other trigeminal nuclei and establish long-range connections to low-level structures of the motor pathway and acetylcholine-releasing nucleus. As described by **Esmaeili et al. (2020)**, acetylcholine is thought to play a role in associating a whisker stimulus with a reward expectation. Through a direct connection from the trigeminal nuclei to PPN, low-level structures of the ascending somatosensory pathways can modulate cholinergic reward signals.

Our findings showed that SPVc inhibits the trigeminal nuclei, PO, SC, VPM, VII, and PPN. Uncategorized connections were found targeting ZI, MOp, and bfd. Connections toward bfd, ZI, PO, VPM, and SPVo were anatomically validated. These findings corroborate previous observations by **Jacquin et al. (1990)**, who demonstrated dense intersubnuclear connections in the rat’s trigeminal complex. Finally, the presence of inhibition in the trigeminal nuclei suggests a significant role for this node in filtering information in these early sensory structures. We could confirm anatomical projections toward bfd, ZI, PO, VPM, and SPVo. A long-range direct projection from SPVc to bfd was shown even though it is sparse. The connection to MOp remains putative for now.

The experiments targeting PSV had a specificity below 75%, therefore, we can only report putative connections from wild-type mice. Projections towards LC, VTA, SI, SPVo, and SPVc were found, but couldn’t be confirmed.

### 3.2 Thalamus

Traditionally, thalamic nuclei of the somatosensory system are considered as relay nuclei that contribute to the spatiotemporal representation of touch (**Ahissar and Arieli, 2001, Azarfar et al., 2018**). It is widely accepted that feed-forward excitatory projections originating from the ventral posteromedial (VPM) and posterior medial (PoM) nuclei, targeting the barrel field cortical columns and septa, respectively, constitute the primary pathway for information flow (**Kleinfeld et al., 1999**). However, our surprising findings argue that these projections do not only include feed-forward excitation but also incorporate feed-forward inhibition. This finding has significant implications, as it reshapes our understanding of intrathalamic communication between the VPM and PoM nuclei.

Long-range connections from VPM originate from both excitatory and inhibitory cells. In addition to directly exciting bfd, Mop (M1), ZI and RT (nRT), VPM sends top-down excitatory projections to PSV. Direct inhibition (Supplemental Figures 7 and 34) of Mop (M1), bfd, ZI, RT (nRT), and PO has been demonstrated to originate from VPM. A connection toward VTA was only found in wild mice. The connections originating in VPM and targeting Mop (M1), ZI, PO, and PSV were anatomically confirmed. The projection to VTA remains putative. These findings suggest that VPM plays a complex role in shaping sensory information flow and sensorimotor transformations within the thalamus.

PO exhibits excitatory connections with RT (nRT), VPM, bfd, and Mop (M1). Uncategorized connections were also identified between ZI. A connection toward SC was seen only in wild mice. The connections targeting VPM were anatomically confirmed, whereas projections toward ZI remain putative.

Inhibitory connections originating from RT (nRT) target ZI, PO, SC and VPM. Connections in wild mice were shown to target VTA, SI, MOp (M1), bfd, and VII. All connections originating from RT (nRT) were results found in an experiment with specificity below 75% specificity, they, consequently, remain putative.

While inhibition of the somatosensory thalamus is generally attributed to teh activity of ZI and RT (nRT) **(Barthó et al., 2002)**, our findings reveal the presence of presynaptic inhibitory neurons in VPM, specifically connecting to PO. Further physiological investigations are warranted to determine whether these projections mediate the inhibition of the paralemniscal pathway by the lemniscal pathway.

### 3.3 Barrel field of the primary somatosensory cortex

The conventional understanding of whisker information processing depicts a bottom-up integration followed by a top-down propagation of neural activity, ultimately leading to the activation of facial muscles. However, our study reveals a more nuanced picture, demonstrating a bidirectional flow of information, with top-down regulation of sensory information from Bfd to the brainstem and thalamus through both excitatory and inhibitory monosynaptic connections.

Bfd extends long-range connections to lower-level structures within the whisker system. Parallel excitatory and inhibitory pathways target MOp (M1), ZI, PO, SC, VPM, PPN, SPVc, SPVi and RT (nRT). Additionally, VTA, VII, PSV and SPVo are excited by bfd, and an uncategorized connection targeting SI was identified.

All new connections originating from bfd were anatomically confirmed, those include PPN, VTA, SI, VII, and SPVo projections, demonstrating top-down modulation of the trigeminal nucleus and facial motor nucleus by Bfd.

### 3.4 Primary Motor cortex

Primary motor cortex extensively targets nuclei across the cerebellar cortex and subcortical regions (see e.g.(**Bisman et al., 2011**, **Jeong et al., 2016**, **Munõz-Castañeda et al., 2021**). Our results show that there are parallel (i.e. excitatory and inhibitory) pathways of projections originating from the primary motor cortex and target bfd, PO, PPN, VTA, SC, VPM, RT (nRT), and ZI from Mop (M1). Monosynaptic projections into these target nuclei have been described before (**Bisman et al., 2011**, **Jeong et al., 2016**, **Munõz-Castañeda et al., 2021**). SI, PSV, SPVo and SPVc also receive excitatory and inhibitory projections. The connection toward SPVc has been anatomically validated.

In addition to the aforementioned projections, excitatory connections were found to target VII and SPVi. Connections toward LC and DR however were witnessed only on wild-type mice. The observations on the parallel feed-forward excitatory and feed-forward inhibitory projections argue that the role of the motor cortical projections might have a differential contribution to neural processing along ascending and descending pathways.

### 3.5 Superior Colliculus

In the map previously described by **Bosman et al., 2011**, the superior colliculus contribution to sensorimotor integration in the 18 nuclei map was limited to connections to ZI, PPN, and VII. Our study revealed that the SC does not only inhibit these nuclei but is also the source of inhibition toward PO, RT and DR. PPN and ZI appear to also receive excitatory connections from the SC. Wild-type connections, however, span the entire sub-network except for TM which doesn’t appear to be targeted by the SC. The newly reported connections toward Mop (M1), RT (nRT) and bfd were anatomically confirmed. Projections toward LC and SPVc remain putative.

### 3.6 Facial motor nucleus

The facial motor nucleus, a key component of the descending motor pathway, is composed of motoneurons and sends commands to the periphery to produce whisker movement (**Ashwell, 1982**).

In this study, we found anatomically confirmed connections originating from VII targeting the trigeminal nuclei and the SC. These structures, being the first nuclei to integrate whisker information in the brain, are also targeting VII (see section 3.1 Trigeminal nuclei). The interplay between VII and the trigeminal nuclei might be part of a low-level loop integrating sensory information and producing whisker movement without requiring activity from higher structures.

More precisely, motor nucleus excites PSV, SPVc and SPVi. Connections that could not be definitively classified as excitatory or inhibitory (i.e. uncategorized) were found to target SPVo, SC, PPN and DR. The connection toward DR was anatomically validated while a connection toward PPN remain putative.

Reciprocal connections between the trigeminal nuclei and VII should be further investigated to understand the role of such low-level loops in the whisker system of the rodent. As VII does not send any inhibitory connections, based on the currently available data, one can assume the purely excitatory nature of the projections to SC. The postsynaptic target of the facial motor nucleus projections into SC is not clear. If this connection targets the sensory area of SC, this suggests the possibility of an efference copy, a signal sent to SC to inform it about the action taken by the whisker and therefore, to adjust its internal representation based on previous movements. The same hypothesis can be made for the trigeminal nuclei interaction.

### 3.7 Zona Incerta

The zona incerta is described as one of the main sources of inhibition in the rodent whisker system (**Venkataraman et al., 2021**). Therefore, unsurprisingly we found inhibitory presynaptic cells projecting to almost all nuclei of the sub-network. These connections were found to target Mop (M1), the thalamic nuclei, PSV, VTA, SI, LC, SC, TM, VII, PPN, RT(nRT) and DR. Wild-type projections connect to SPVo and SPVi New connections targeting bfd, PSV, SI, SPVo, VPM, VII, SPVc and RT (nRT) were anatomically confirmed. A connection toward SPVi, on the other hand, remains putative.

In line with most studies targeting ZI reporting GABAergic connections (**Ficalora and Mize, 1989**, **Nicolelis et al., 1992**, **Nicolelis et al., 1990**), we didn’t find any excitatory presynaptic cells in ZI projecting the sensorimotor system. Furthermore, the extensive connectome of ZI spanning the somatosensory pathway and in particular low level structures such as the trigeminal nuclei and VII hint at a prominent role played by ZI in low-level sensorimotor integration.

### 3.8 Neuromodulatory projections

The VTA has a role in learning the association of a whisker stimulus with a potential reward (**Esmaeili et al., 2020**). Our results revealed that the VTA’s contribution to the whisker system’s function is not necessarily solely mediated by dopaminergic connections; excitatory presynaptic neurons in the VTA also target other nuclei. The VTA sends excitatory connections to TM, DR, SI, MOp (M1), bfd, ZI, VPM and RT (nRT). Projections toward SPVo and SPVi were identified, but the transgenic line showing those connections couldn’t be categorized as either excitatory or inhibitory. Each of the new connections reported herein towards bfd, SPVo, TM, VPM, RT and SPVi remain putative.

Cholinergic connections have been hypothesized to play a role in transforming whisker sensations into a motor action (e.g. licking behavior (**Esmaeili et al., 2020**)). Consequently, it is not surprising to see connections originating from SI that target bfd and Mop (M1). However, as for VTA, we observed a large number of connections, mainly inhibitory, targeting other neuromodulator-releasing nuclei. This shows that SI can regulate the activity of other nuclei in the neuromodulatory system. In particular, SI inhibits PPN, another cholinergic nucleus, TM, VTA, as well as MOp (M1), bfd, ZI, RT (nRT), and DR. One connection with non-specific presynaptic cells was unveiled toward SC. New connections to TM, and PPN were anatomically confirmed. The connection toward SC remains putative.

Excitatory connections from DR were found to target PPN, VTA, SI, MOp(M1), bfd, ZI, RT, SC, VII. Uncategorized connections towards PO, PSV, SPVo, LC, TM, VPM, SPVc and SPVi were identified. The connections targeting LC and SI were anatomically confirmed. Projections toward TM, ZI, PO, RT, PSV, SPVo, SPVi and SPVc remain putative.

Inhibitory connections originating within PPN were identified to project to MOp (M1), ZI, PO, bfd, VTA, SI, SC, VII, RT (nRT), and DR. Connections with uncategorized presynaptic cells targets PSV, SPVo, LC, TM, VPM, SPVc and SPVi. PPN targets DR, SI, and RT (anatomically confirmed). Putative connections toward LC, TM, MOp, and bfd were discovered but couldn’t be validated.

### 3.9 Density distribution

To elucidate connectivity patterns within and between structures in the sensorimotor pathway, we conducted an analysis of the density, defined as the percentage of outgoing connections originating from a single source, for each of the 57 nuclei. This examination revealed distinctive connectivity profiles, with the cortex displaying sparse outgoing connections to a multitude of nuclei. The top 6 connections from MOp, MOs, and bfd accounted for at most 78% of the outgoing density (**Figure 5**), emphasizing the distributed nature of cortical inputs. Contrastingly, the Thalamus exhibited a more specialized pattern, with the top 4 of their outgoing projections accounting for over 94% of all of their projections (**Figure 5**).

**Fig. 5.**
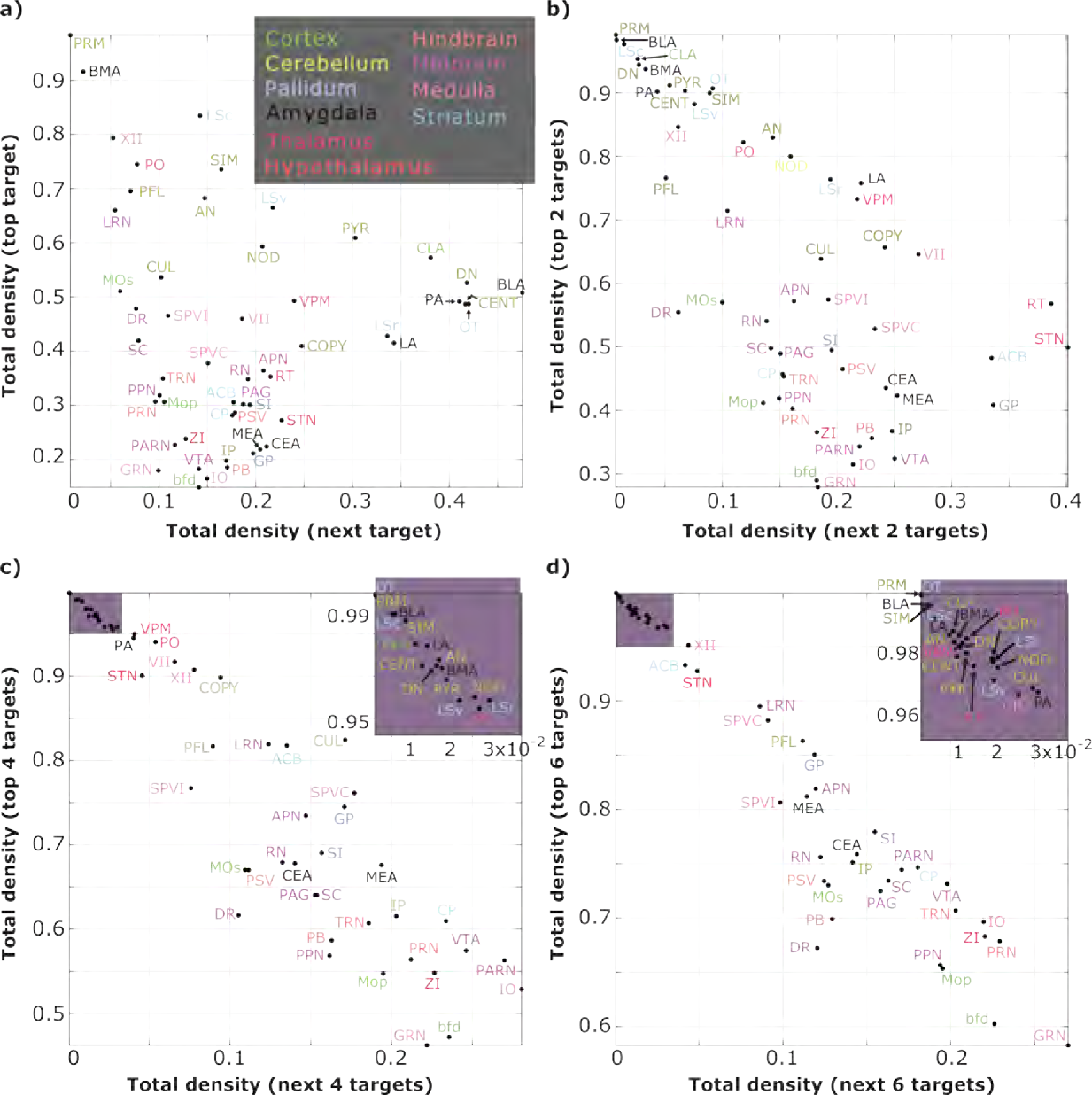
Relative density of outgoing connections. Connections were ordered by the density of outgoing connections and plotted against the *x* next densest connections for comparison. This shows the skewness of the distribution. The color scale represents the anatomical structure to which the nuclei belong. **a)** The density of incoming connections as a function of the density of the next incoming connection (in order of magnitude) **b)**Same as a), but now for the next 2 connections. **c)** Same as a), but now for the next 4 connections. **d)** Same as a), but now for the next 6 connections. Onsets in c) and d) are a zoom-in on the nodes on the top-left corner. FN, LC, TM and SPVo were not included as no connections originating from those nuclei were found, due to a lack of experiments targeting these nodes.

When focusing on the ascending somatosensory pathway, a discernible pattern emerges. The trigeminal nuclei, serving as the initial nodes for whisker information, project sparsely to a diverse array of nuclei. Their top 6 projections targets explain approximately 74% of the outgoing projections (**Figure 5**). Subsequently, the pathway narrows as it targets the thalamus, projecting to fewer post-synaptic nuclei, with 94% of their outgoing density explained by only 4 projections (**Figure 5**). Finally, the cortical information flow displays a divergent pattern, with only 60% outgoing density distributed among 6 nuclei (**Figure 5**).

This observed pattern along the ascending somatosensory pathway underscores a dynamic process: the brainstem initially projects broadly, supporting a divergent network formation, followed by a sharp reduction of the target nuclei of the thalamus. The pathway then broadens again as cortical projections target a diverse set of nodes. This highlights the thalamus’s specific role as a crucial node for routing sensory information. Coupled with inhibitory processes, the thalamus emerges as a key filter, emphasizing its pivotal role in shaping and modulating sensory inputs

### 3.10 Network analysis

To assess the role of neuromodulator-releasing nuclei in shaping the network structure and to investigate their potential facilitation of communication within the network, we conducted a permutation test with 10,000 permutations on the 57-nuclei adjacency matrix (**Supplemental Figure 2**). Comparisons between the network with and without neuromodulator-releasing nuclei revealed no discernible differences in terms of the distribution of betweenness centrality, degree centrality, nodal clustering, nodal efficiency, or local nodal efficiency. However, a notable distinction emerged in the comparison of nodal shortest path lengths. Specifically, the network without neuromodulator-releasing nuclei exhibited a significantly higher mean value (0.8969 ±0.1093 std) compared to the network with neuromodulators (0.8577 ±0.1063 std; p-value = 0.0011). These findings underscore the nuanced impact of neuromodulator-releasing nuclei on the structural dynamics of the network, particularly in terms of nodal shortest path lengths.

To determine if the network presented small-world properties, we compared the clustering coefficient and the shortest path between a structured graph (ER) and the 57×57 network resulting from our analysis of the Allen Brain Database. This analysis revealed a significantly higher average shortest path for our network (mean ER = 0.7159 ±0.0447 std; mean summary graph = 0.8577 ±0.1063 std; p-value = 9.999×10^−5^) and a significantly higher clustering coefficient for our network (mean ER = 0.1241 ±0.288 std; mean summary graph = 0.1726 ±0.1107 std; p-value = 0.0015). Given that our network presents a higher shortest path than an Erdos-Renyi graph (see Materials and Methods), we conclude that the studied network doesn’t present a small-world structure.

Next, we computed the degree centrality of each nucleus of the network (see section 2.5). This analysis revealed that the majority of connections targeted and originated from the midbrain, which represents 21% of all connections. In particular, DR, PPN, SC, PAG, and RN appear to be hubs. The Medulla, while representing 11% of all connections appears to have evenly distributed connections along all its nuclei. In the hindbrain, PB is the only hub despite the hindbrain representing 11% of total connections. The striatum appears to concentrate 9% of connection mainly targeting and originating from the only hub of this structure CP. The cortex represents only 10% of all connections but MOs, MOp, and bfd appear to be hubs, only CLA doesn’t satisfy the criterion. No hubs were identified in either the Cerebellum (10% of connections) the Amygdala (8% of connections) or the Thalamus (5% of connections). In the Hypothalamus (6% of connections), ZI is a hub and the only node that we considered from the Pallidum (5% of connections), GP, is considered a hub. Amongst the nodes, VTA, GRN, PRN, SI, CEA, and IP appear slightly below the threshold to be considered hub. More experiments performed on these nuclei might increase their degree centrality (**Figure 6**).

**Fig. 6.**
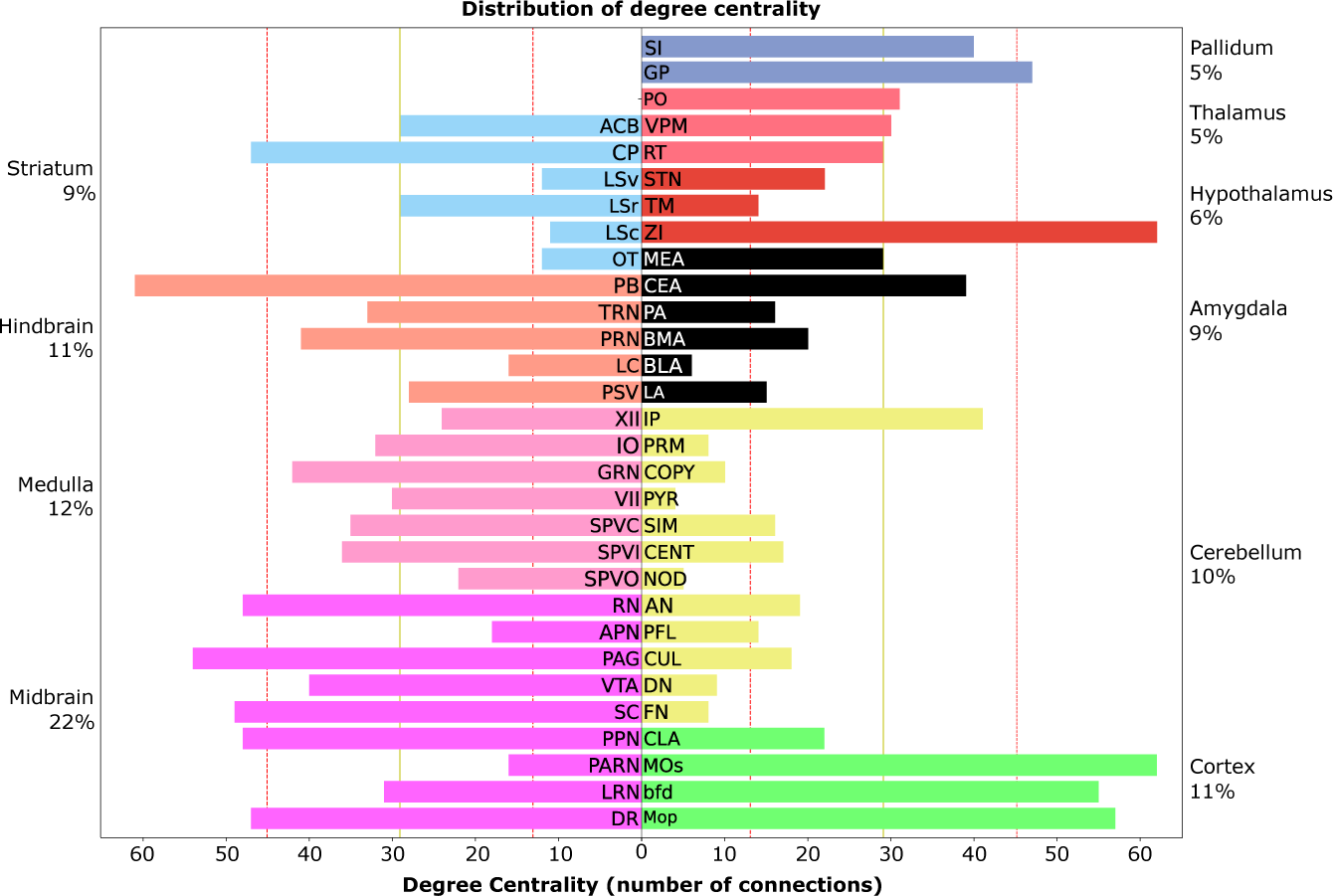
Degree centrality of the 57-nodes network. The yellow line represents the mean of the distribution and the red dotted lines represent the mean ± the standard deviation. We consider every nodes above mean+standard deviation to be a hub.

## 4 Discussion

We extended the established connectome of the mouse whisker system (**Bosman et al., 2011**) by incorporating 42 additional connections (**Figure 7**). We substantiated the presence of these connections through anatomical evidence (**Supplementary Material - Fig.6 to Fig.35**). Within this framework, we identified cell-type-specific connections and hubs, revealing intricate connectivity patterns along the ascending somatosensory pathway.

**Fig. 7.**
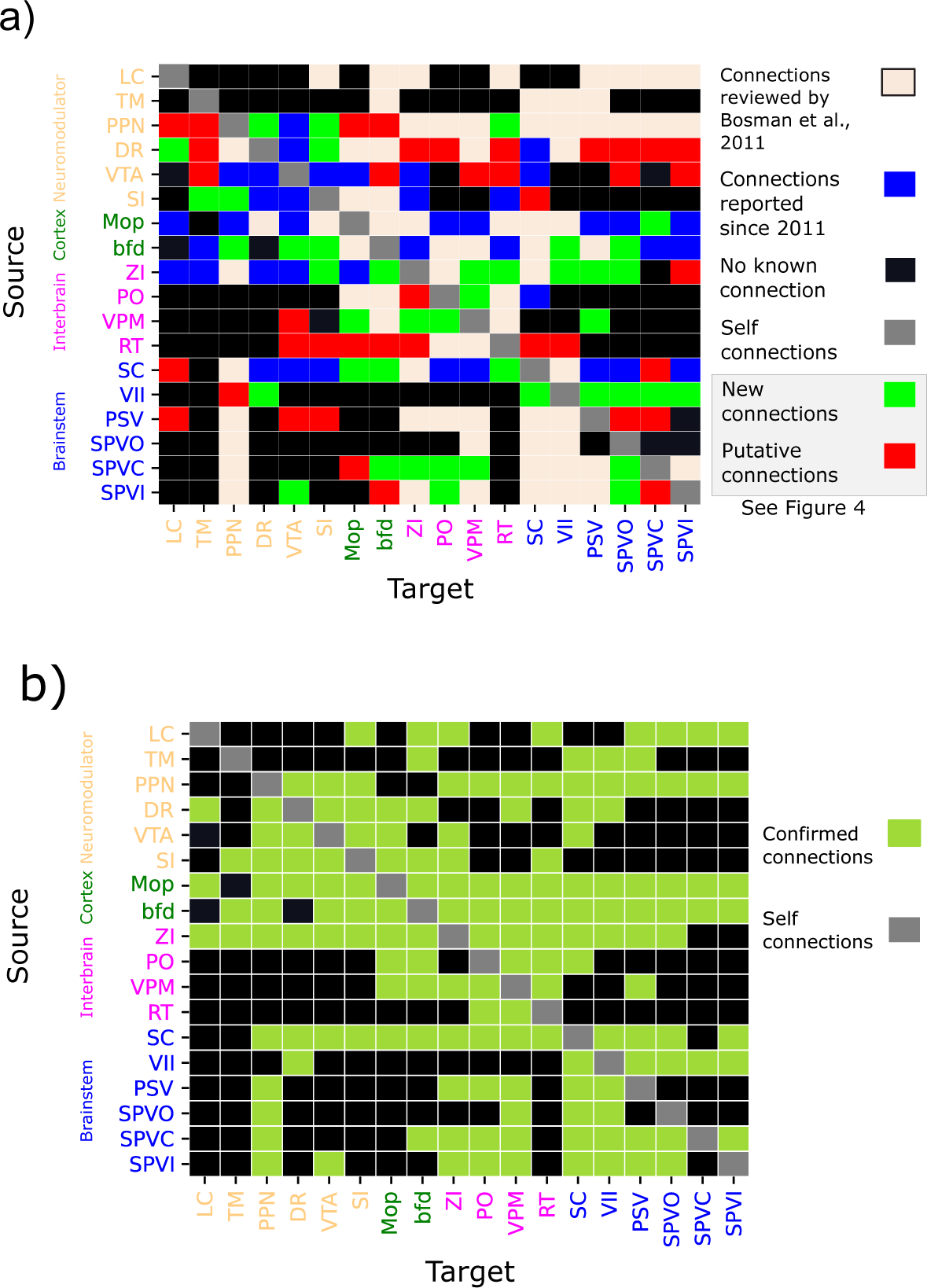
The sensorimotor connectivity map of the mouse whisker system. Summary of data from all animal lines combined. **a)** A total of 157 connections were found and confirmed either by previous work or through anatomical validation. See Figure 4 for new and putative connections only. **b)** A binary connectivity matrix that summarizes the pairwise connectivity in the network with only previously reported or anatomically confirmed connections.

We reported a total of 157 connections, constituting 48.4% of all possible connections within this 18×18 subnetwork. Notably, 42 of these connections were not previously described in the literature, to the best of our knowledge. The 18 nuclei were chosen because they were experimentally shown to contribute to sensorimotor integration, which might explain the high level of connectivity described herein. These findings suggest that even if some nuclei were ‘cut-off’ from the network, information flow could persist, underscoring the robustness of the network organization. Conversely, the analysis of shortest-path length highlights the importance of neuromodulator-releasing nuclei in facilitating faster communication between network nodes. The average shortest path length significantly increases in the absence of neuromodulatory nuclei.

The connectivity pattern of the ascending somatosensory pathway follows an hourglass shape, with divergent brainstem projections, followed by convergence of thalamic projections into only a few targets in the cortex. Once the sensory information reaches the cortex, the divergence of projections once again allows the distribution of the sensory information across a large number of nuclei. This is further supported by the results shown from our analysis of outgoing densities (**Figure 5**, also see **Figure 6**). In fact, this analysis demonstrates that the trigeminal nuclei are nodes that distribute information beyond the sensory thalamus. Meanwhile, the narrowing of the pathway at the level of the thalamus through denser and fewer connections, together with the presence of inhibition in this structure, might hint at a filtering role of the somatosensory thalamus, as it appears to be highly specialized given the few strong connections originating from this structure (**Figure 5**). Further widening of the projections in the cortex points to possible parallel processing of sensory information along the downstream nuclei.

### 4.1 Prominent inhibitory projections in every stage of the ascending pathway

Inhibition is a pervasive phenomenon at every level in the whisker system. Our study reveals inhibitory contributions from nodes not previously identified as sources of inhibition. A notable example is the inhibitory connection from VPM to PO, as illustrated in the Ppp1r17-Cre-NL1 animal line (**Supplemental Figure 6**). This discovery challenges the notion that RT (nRT) is the sole source of inhibition in the somatosensory thalamus. The inhibitory pathway from VPM may even play a role in disinhibition, given its targeting of RT (nRT) and ZI, which, in turn, influence the thalamic nuclei of the ascending somatosensory pathway (**Figure 3**). This suggests a nuanced filtering or processing role for VPM beyond its traditional role as a relay station.

Inhibition is not confined to higher levels; it is also observed in the trigeminal nuclei, which send inhibitory connections (**Figure 3**). While these connections primarily target other trigeminal nuclei, especially VII, they may play a crucial role in modulating whisker movement amplitude following contact with an object during free whisking. Descending projecting connections from higher-level structures could then further modulate whisking behavior during active exploration. This low-level loop in the whisker system underscores the ability of trigeminal nuclei to reduce the whisking amplitude upon contact with an object. Furthermore, ascending inhibitory connections from the somatosensory brainstem (trigeminal nuclei) target VPM and PPN, suggesting a multifaceted role for the trigeminal nuclei beyond a simple relay station in the whisker system. In conclusion, our analysis points to extensive and intricate processing that involves both excitatory and inhibitory projections in the early phases of the somatosensory pathway, as it was previously described in invertebrates for sensorimotor control (**Hughes and Celikel, 2019**)

### 4.2 Top-down modulation of sensorimotor integration: a primer

While the structures under consideration play a role in sensorimotor integration, it is plausible that some of the identified connections may not directly contribute to sensorimotor processing. Notably, certain structures (**Supplementary Material - Table 2**), such as neuromodulator-releasing nuclei, were targeted by only a few, if any, experiments. These connections warrant further investigation to better understand their functional significance. However, to ensure the robustness of our findings and mitigate potential biases from false positives or negatives, we confirmed the connections through anatomical validation (**Figure 4**).

The Pedunculopontine Nucleus (PPN) is the most interconnected node among neuromodulatory nuclei, it appears to send connections to every node of the somatosensory system, with the exception of the cortex. Interestingly, the somatosensory thalamus doesn’t appear to project to PPN. The Dorsal Raphe nucleus (DR) stands out amongst the neuromodulator-releasing nuclei as a node that doesn’t receive direct input from the sensory pathway. LC seems to target mainly structures in the low-level sensory system. SI doesn’t appear to have a role in the regulation of activity in low-level structures, as no connections toward the trigeminal nuclei or VII could be confirmed. Notably, the neuromodulator-releasing nuclei display a high degree of interconnectedness, allowing for mutual modulation.

Top-down regulation is a prominent feature in the somatosensory system, occurring both between the barrel field cortex (bfd) and the thalamus and between the thalamus and the trigeminal nuclei. Inhibitory projections, particularly originating in bfd and targeting VPM, RT (nRT), and PO, run in parallel to excitatory connections, allowing for effective top-down modulation of somatosensory thalamic activity. Additionally, a direct top-down connection from VPM to PSV highlights the regulatory influence of the lemniscal pathway.

### Shortcomings of the experimental approach

We used anatomical data collected through serial 2-photon tomography, made available by the Allen Brain Institute, to confirm targets and the source of the monosynaptic connection. This method allows visualization of the expressions of genetically encoded fluorescent proteins in anatomically targeted regions using cell-type-specific or non-specific promoters in various animal lines. However, it’s important to note that the spatial resolution of this technique is not sufficient to resolve individual synapses, so it is not always possible to reliably distinguish en-passant projections from actual terminals. Additionally, the sensorimotor circuit outlined in our analysis is derived from anterograde tracing experiments. To further refine our understanding, trans-synaptic tracing of each discussed connection would be beneficial, enabling differentiation between en-passant fibers and terminals and providing cell-type information for the pre- and post-synaptic neurons.

Our results are based on the analysis of data from 732 different experiments. To anatomically confirm the presence of the connections mentioned in our results section and depicted in our summary map (**Figure 7**), we employed two-photon tomography imaging in brain slices. However, challenges arose during the alignment of the normalized brain atlas to each image due to variations in cut angles. Furthermore, tissue alterations caused by cuts, such as compression, and potential tissue shrinkage induced by the fixation solution injection added complexity to the alignment process. The inherent discretization of imaging data, in contrast to the continuous nature of data analysis, also presented challenges in achieving accurate alignment between the normalized brain and imaging data.

Finally, our analysis relies entirely on the Allen Brain database annotation and only integrates structures available in the database. For instance, a structure such as the nucleus basalis of Meynert is considered in this study as a subpart of SI as it is not presented separately in the database. We are, therefore, limited to the granularity of analysis allowed by the Allen Brain database, thus, some of the structures presented in previous works (**Bosman et al., 2011**) were not included in this study.

### Are mice small rats?

In their prior study, Bosman et al. (**2011**) considered connections from various rodent species, encompassing both rats and mice, assuming uniformity across the whisker systems of different rodent subspecies. However, a review by Krubitzer et al. (**Krubitzer et al., 2011**) highlighted significant anatomical differences in rodents, particularly between rats and mice. Despite these differences, the two species are often treated interchangeably in anatomical structure sizes or positions, raising questions about the relevance of employing an average model for the broader category of rodents, where connections are constrained by brain anatomy.

The connections we demonstrate in this study are based on experiments conducted specifically on mice, avoiding the potential constraint imposed by assuming a uniform rodent model. Nevertheless, the original connections we build upon were not observed in studies that differentiated between rats and mice. To thoroughly assess the similarity between the rat and mouse connectomes, further translational anatomical studies will be necessary.

## Conclusions

Leveraging the Allen Brain database, we identified 42 new connections that were anatomically validated and 40 putative connections, categorizing their nature and enhancing our understanding of the subnetwork’s connectome. Our analysis emphasizes the role of both low-level structures and neuromodulator-releasing nuclei, revealing the widespread occurrence of inhibition starting from the earliest stages of information processing. While this study provides a clearer picture of how the sensorimotor pathway integrates whisker information as well as the contributions of the top-down and neuromodulatory circuits, further investigations, including functional studies and testing with additional transgenic lines, are necessary to elucidate the specific roles of each connection in sensorimotor integration. The road to a comprehensive understanding of the rodent whisker system’s sensorimotor pathway requires more extensive circuit analysis, including trans-synaptic tracing experiments, particularly with higher granularity on low-level structures and neuromodulator-releasing nuclei, to pave the way for the development of a biologically inspired computational model of the entire network.

## List of abbreviations

TG: Trigeminal Ganglion
PSV: Principal sensory nucleus
SPVO, SPVI, SPVC: spinal nucleus of the trigeminal oralis, interpolaris, caudalis
VPM: ventral posteromedial nucleus
PO: posterior complex of the thalamus
RT (nRT): Reticular Nucleus of the thalamus
bfd: barrel field of the primary somatosensory cortex
MOp (M1): primary motor cortex
SC: Superior Colliculus
VII: Facial Motor nucleus
ZI: Zona Incerta
SI: Substantia Innominata
DR: Dorsal raphe nucleus
LC: Locus coeruleus
PPN: Pedunculopontine nucleus
VTA: Ventral tegmental area
TM: Tuberomammillary nucleus

## Acknowledgment

This project has received funding from the European Union’s Horizon 2020 research and innovation program under the Marie Sk-lodowska-Curie grant agreement No 860949.

## Supplementary Materials

**Supplemental Table 1:**
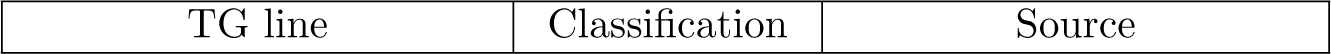
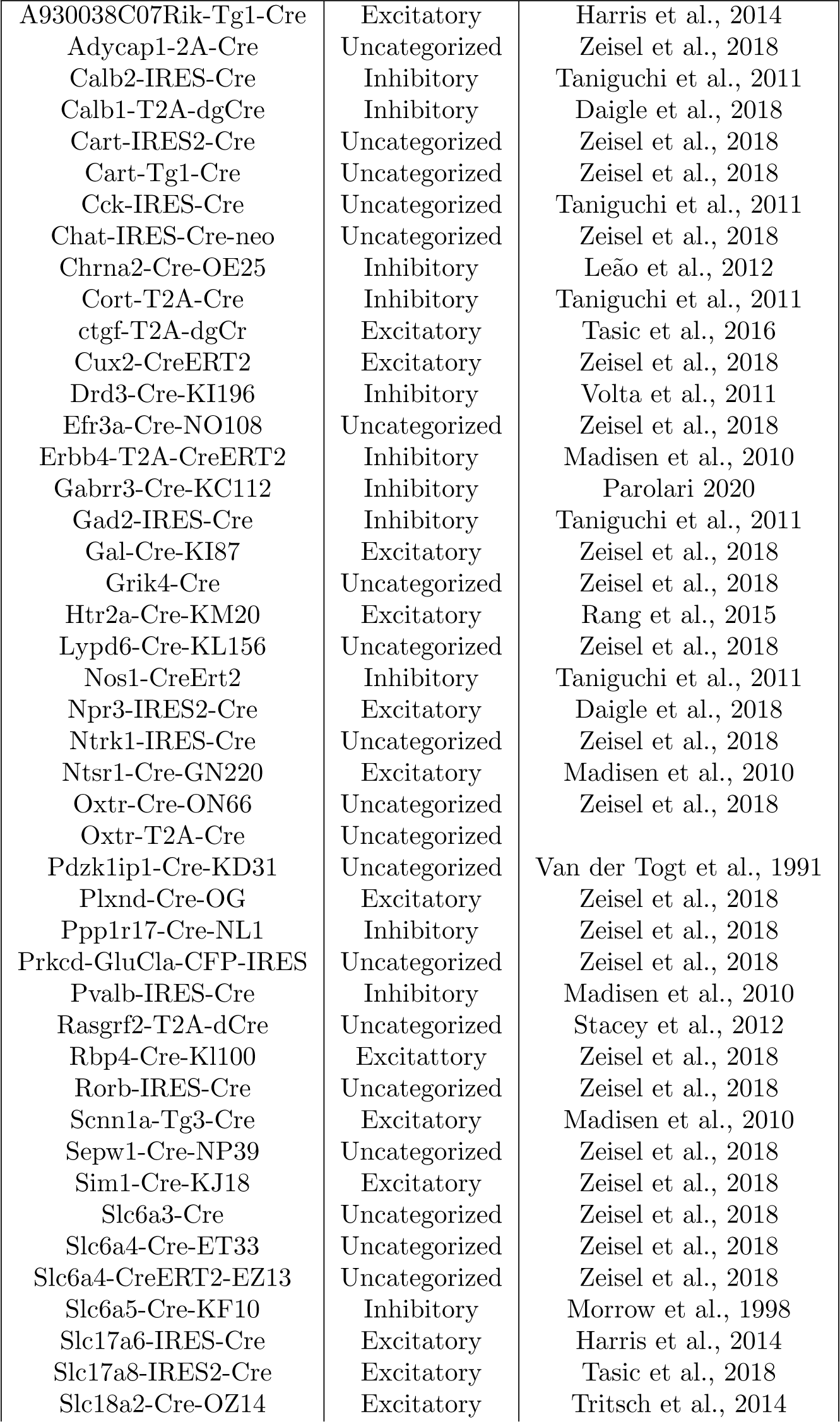
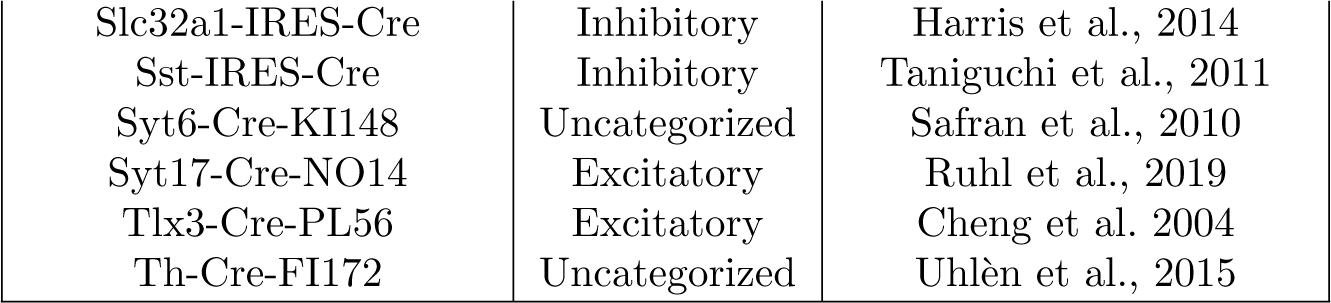
Type of neuron marked by2e0a1c5h transgenic line.

**Supplemental Table 2:** For each structure, this table presents the number of experiments ran (transgenic and wild), the number of transgenic lines tested, the number of transgenic lines showing marked neurons in the considered structures and the number of transgenic lines showing a connection.

| structure | wild type<br>experi-<br>ments | transgenic<br>line<br>experi-<br>ments | transgenic<br>lines | expressed<br>in num-<br>ber of<br>struc-<br>tures | transgenic<br>line<br>showing<br>connec-<br>tions |
| --- | --- | --- | --- | --- | --- |
| MOp | 8 | 21 | 15 | 13 | 12 |
| ZI | 4 | 15 | 13 | 11 | 11 |
| PO | 3 | 6 | 3 | 3 | 3 |
| bfd | 6 | 40 | 29 | 21 | 16 |
| PSV | 2 | 1 | 2 | 1 | 1 |
| VTA | 2 | 12 | 12 | 9 | 8 |
| SI | 7 | 10 | 8 | 7 | 7 |
| SPVO | 1 | 0 | 0 | 0 | 0 |
| LC | 0 | 0 | 0 | 0 | 0 |
| SC | 7 | 14 | 9 | 7 | 7 |
| TM | 0 | 1 | 1 | 0 | 0 |
| VPM | 3 | 6 | 6 | 5 | 5 |
| VII | 2 | 5 | 4 | 3 | 3 |
| PPN | 0 | 3 | 3 | 2 | 2 |
| SPVC | 3 | 6 | 6 | 5 | 5 |
| SPVI | 2 | 2 | 1 | 1 | 1 |
| RT | 2 | 11 | 7 | 7 | 7 |
| DR | 0 | 9 | 5 | 4 | 4 |

**Supplemental Fig. 1.**
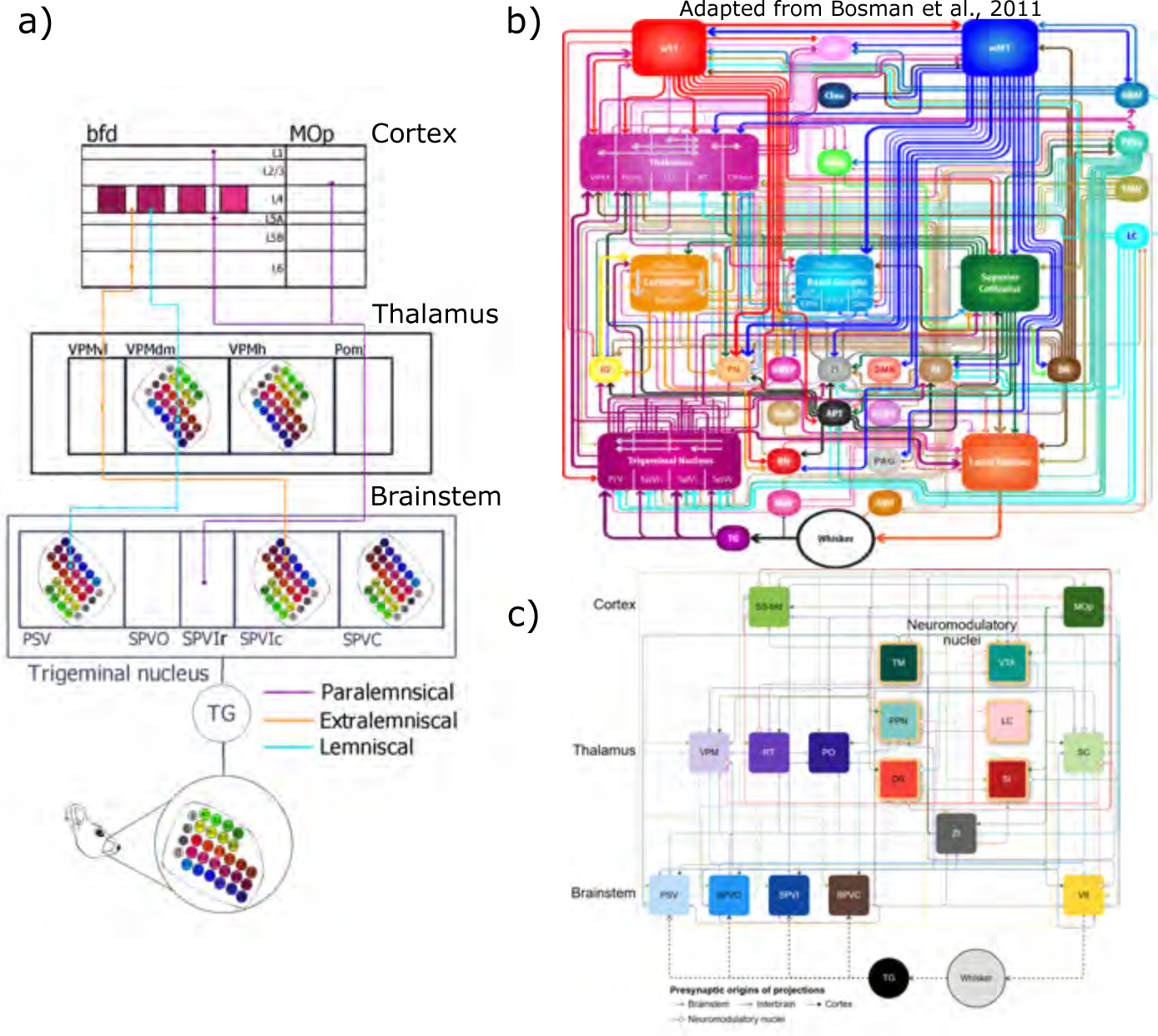
a) The whisker pad is organized in a grid-like structure. Each whisker is represented in SPVI, PSV, VPM, and bfd by an agglomerate of neurons organized in a barrel-like shape, respecting the coordinates of the whisker on the whisker pad with rotations. The information from the whisker enters the system through the trigeminal ganglion, projecting to the trigeminal nuclei. Here the information flow is divided into three major pathways: 1) the lemniscal pathway (blue), originating from the barrelettes of PSV, ascending to the barreloids of VPMdm and ending in the barrels of the layer 4 of the somatosensory pathway; 2) the extralemniscal pathway (orange), taking its origin in inter-barrelette neurons of SPVI, connecting to VPMvl and targeting the primary somatosensory cortex; and 3) the paralemniscal pathway (purple), taking its source in SPVI, targeting PO, which then targets MOp and bfd. b) The connectivity map presented by Bosman et al. (2011).The 18 nuclei that we focus in our study present 72 connections in the previous map. c) New connectome composed of 157 connections.

**Supplemental Fig. 2.**
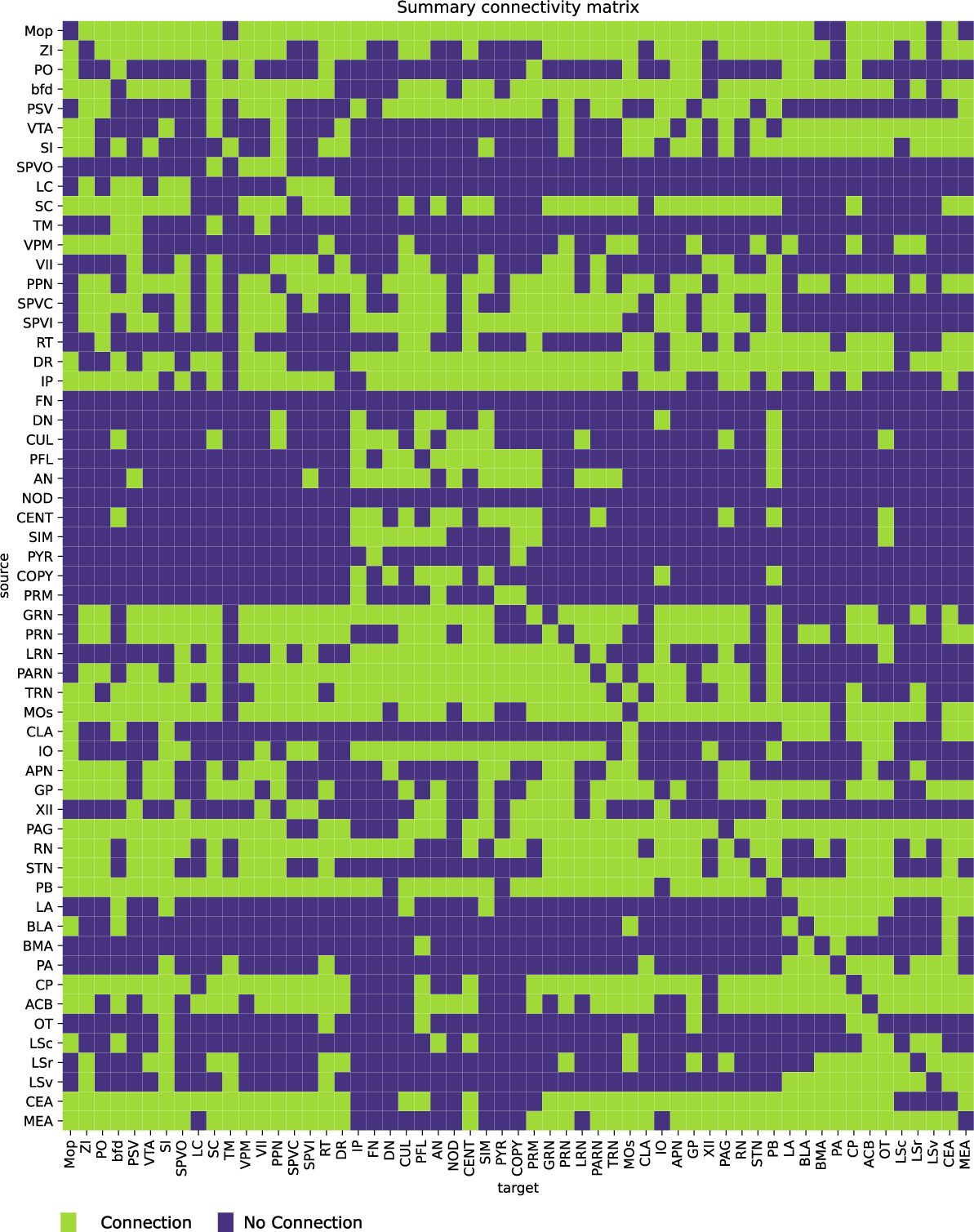
Summary of connection between 57 nuclei. **MOp**: Primary Motor Cortex, **Zi**: Zona Incerta, **PO**: Posterior complex of the thalamus, **bfd**: Barrel field of the primary somatosensory cortex, **PSV**: nucleus principalis of the trigeminal nucleus, **VTA**: Ventral tegmental area, **SI**: Substantia Innominata, **SPVO**: Spinal nucleus of the trigeminal nucleus oral part, **LC**: Locus coeruleus, **SC**: Superior Colliculus, **TM** : Tuberommaimillary Nucleus, **VPM**: Ventraposteromedial nucleus of the thalamus, **VII**: Facial motor nucleus, **PPN**: Pedunculopontine nucleus, **SPVC**: Spinal nucleus of the trigeminal nucleus caudal part, **SPVI**: Spinal nucleus of the trigeminal nucleus interoplaris part, **RT**: Reticular nucleus of the thalamus, **DR**: Dorsal Raphe nucleus, **IP**: Interposed nucleus, **FN**: Fastigial nucleus, **DN**: Dentate nucleus, **CUL**: Culmen, **PFL**: Paraflocculus, **AN**: Ansiform lobule, **NOD**: Nodulus(X), **CENT**: Central lobule, **SIM**: Simple lobule, **PYR**: Pyramus(VIII), **COPY**: Copula pyramidis, **PRM**: Paramedian lobule, **GRN**: Gigantocellular reticular nucleus, **PRN**: Pontine reticular nucleus, **LRN**: Lateral reticular nucleus, **PARN**: Parvicellular reticular nucleus, **TRN**: Tegmental reticular nucleus, **MOs**: Secondary motor area, **CLA** : Claustrum, **IO**: Inferior olivary complex, **APN**: Anterior pretectal nucleus, **GP**: Globus Pallidus, **XII**: Hypoglossal nucleus, **PAG**: Periaqueductal gray, **RN**: Red nucleus, **STN**: Subthalamic nucleus, **PB**: Parabrachial nucleus, **LA**: Lateral amygdalar nucleus, **BLA**: Basolateral amygdalar nucleus, **BMA**: Basomedial amygdalar nucleus, **PA**: Posterior amygdalar nucleus, **CP**: Caudoputamen, **ACB**: nucleus accumbens, **OT**: Olfactory tubercle, **LSc**: Lateral Septal nucleus caudal part, **LSr**: Lateral Septal nucleus rostral part, **LSv**: Lateral Septal nucleus ventral part, **CEA**: Central amygdalar nucleus, **MEA**: Medial amygdalar nucleus

**Supplemental Fig. 3.**
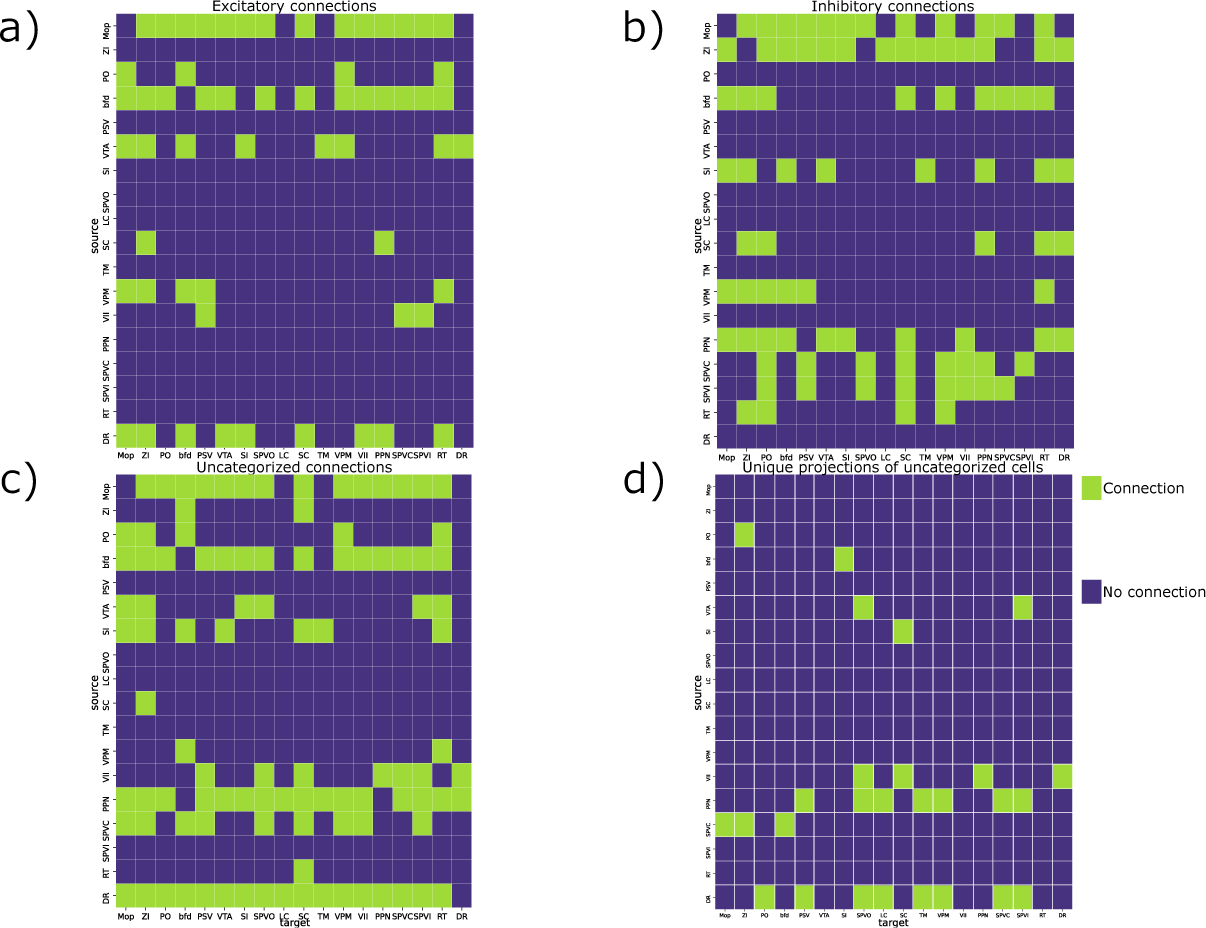
**a)** The excitatory connectivity matrix contains 58 connections. **b)** The inhibitory connectivity matrix contains 84 connections. **c)** 101 connections with uncategorized presynaptic cells were unveiled. **d)**27 connections were shown only by uncategorized transgenic-line

**Supplemental Fig. 4.**
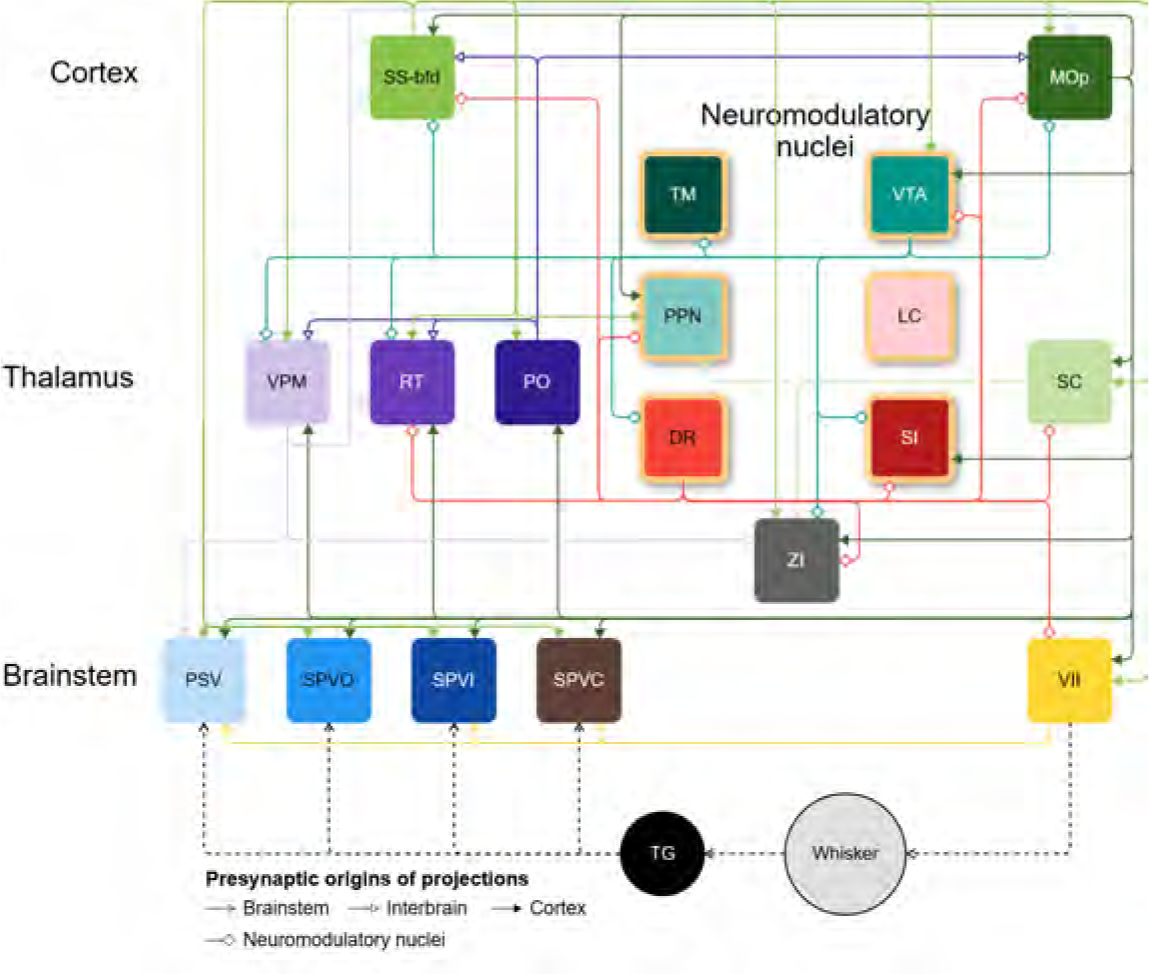
The excitatory connectivity map contains 61 connections. The dashed line represents the flow of information outside of the central nervous system. The neuromodulatory structures are represented with yellow borders.

**Supplemental Fig. 5.**
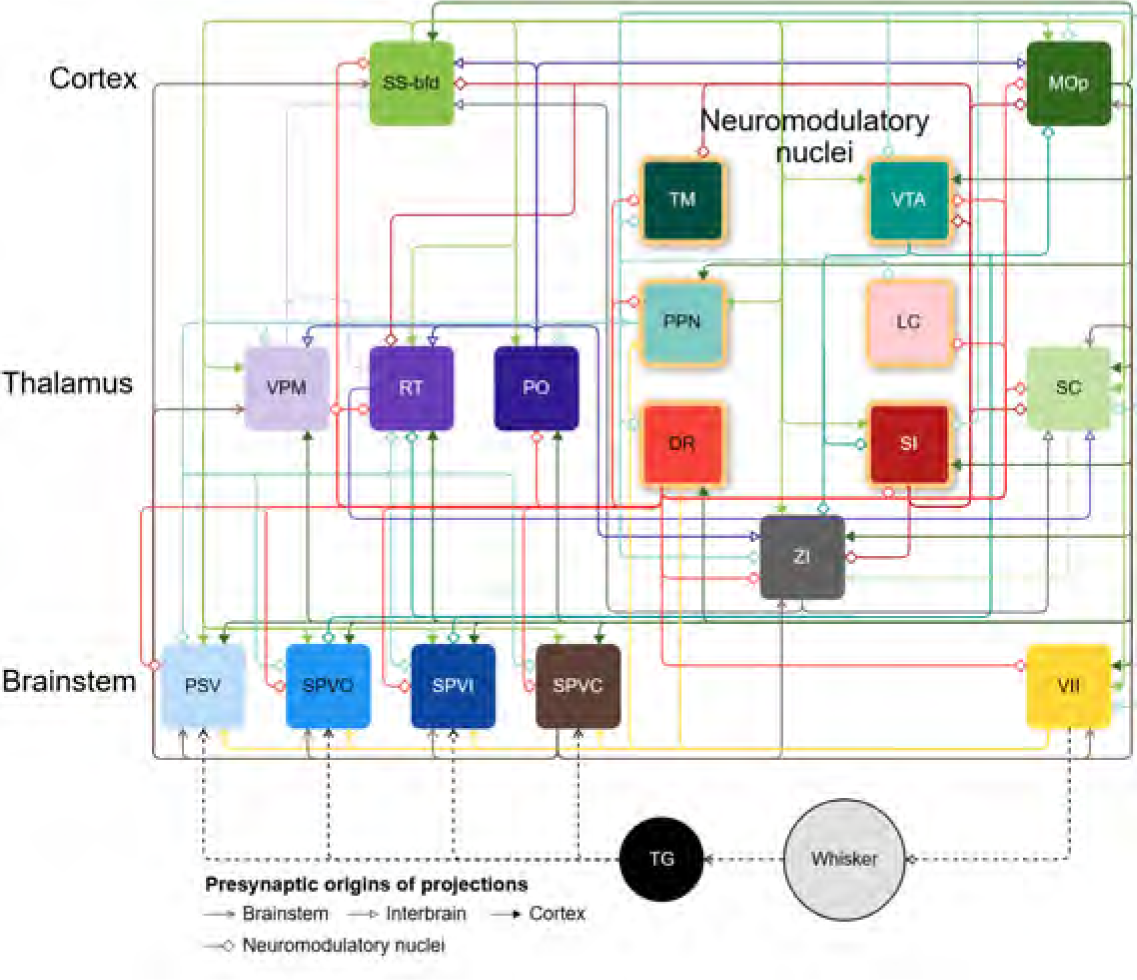
95 connections were revealed by uncategorized transgenic lines. The dashed line represents the flow of information outside of the central nervous system. The neuromodulatory structures are represented with yellow borders.

**Supplemental Fig. 6.**
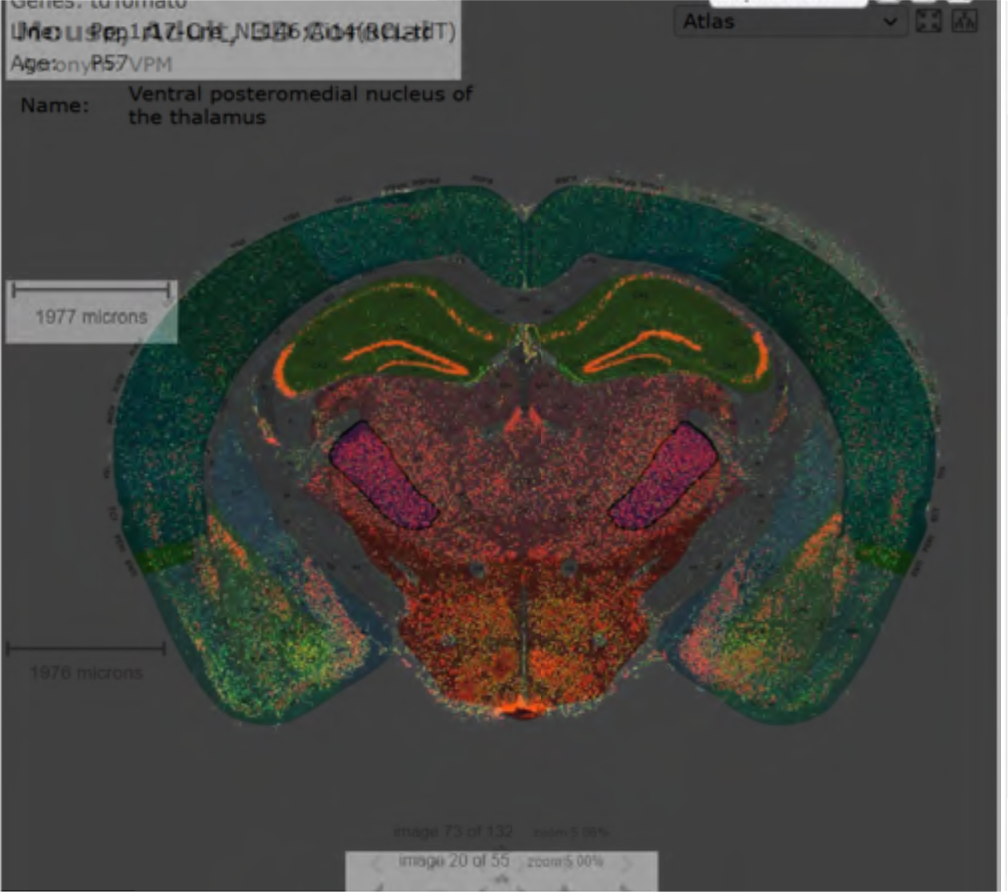
Expression in VPM of the transgenic lines PPP1r17 (inhibitory)

**Supplemental Fig. 7.**
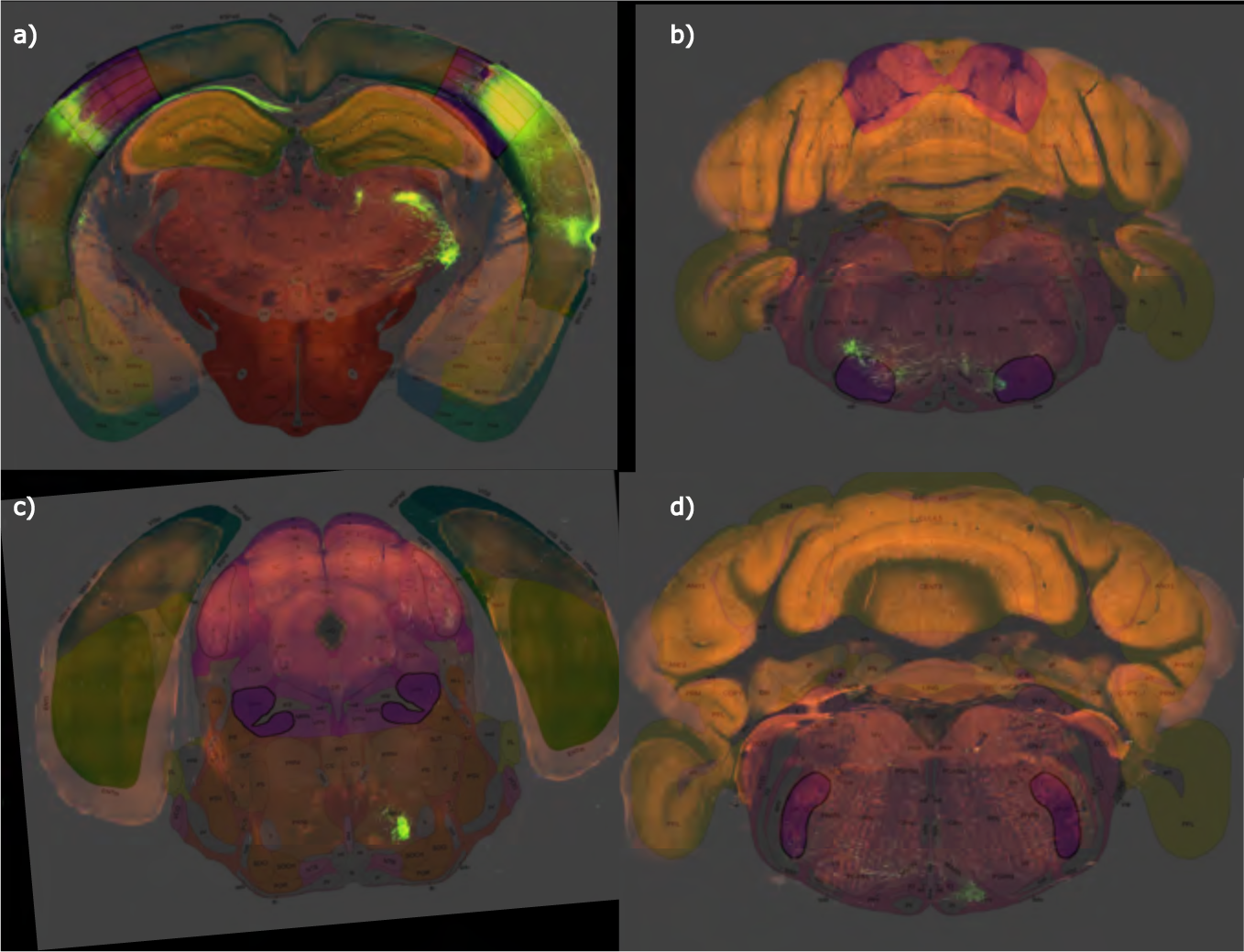
**a)** Injection structure bfd to **b)** VII, **c)** PPN, and **d)** SPVo. Experiment number 112951804, wild-type. The shortcomings of experimental procedures are addressed in the discussion.

**Supplemental Fig. 8.**
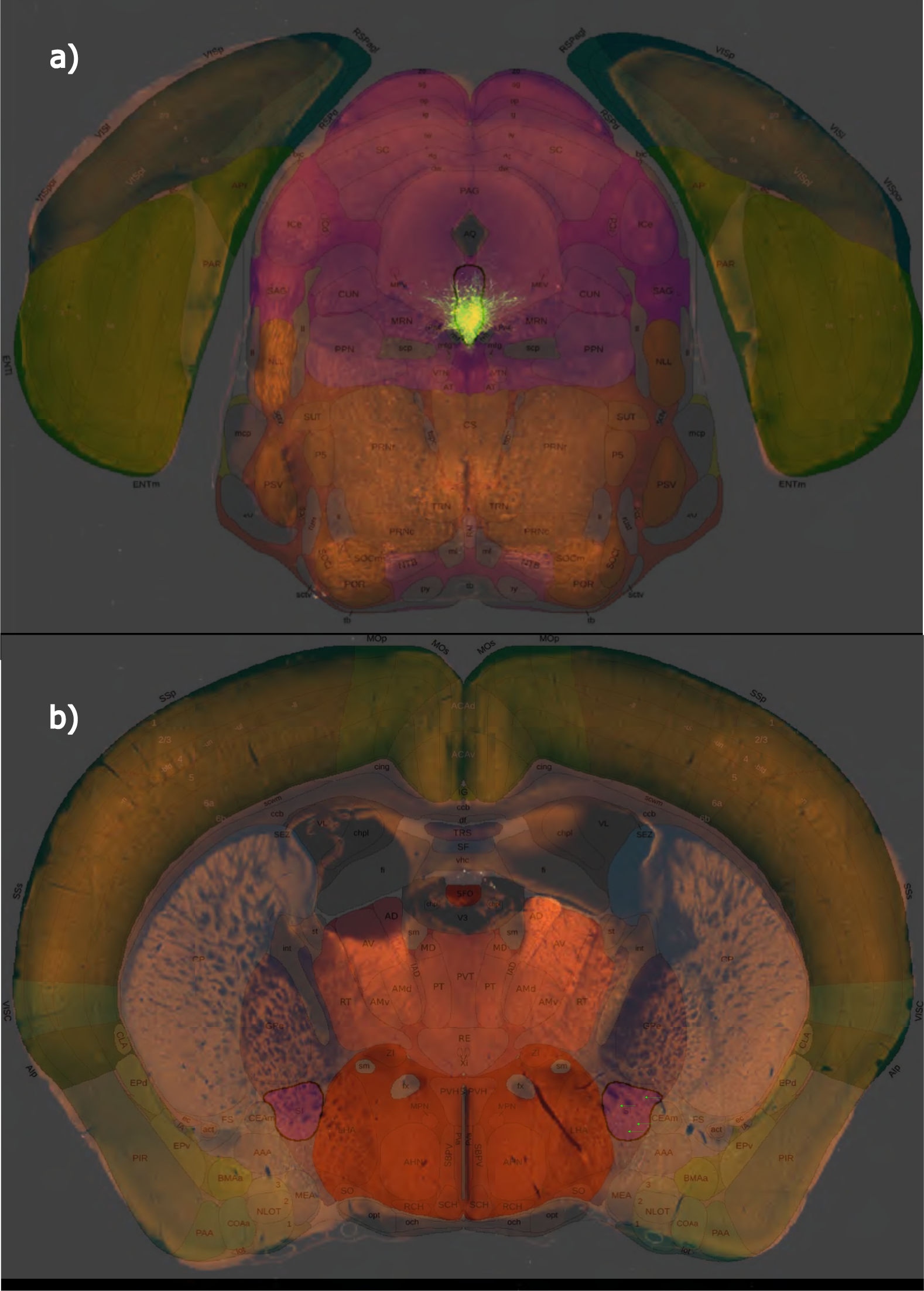
**a)** Injection structure DR to **b)** SI. Experiment number 582609848, line Esr2-IRES2-Cre. The shortcomings of experimental procedures are addressed in the discussion.

**Supplemental Fig. 9.**
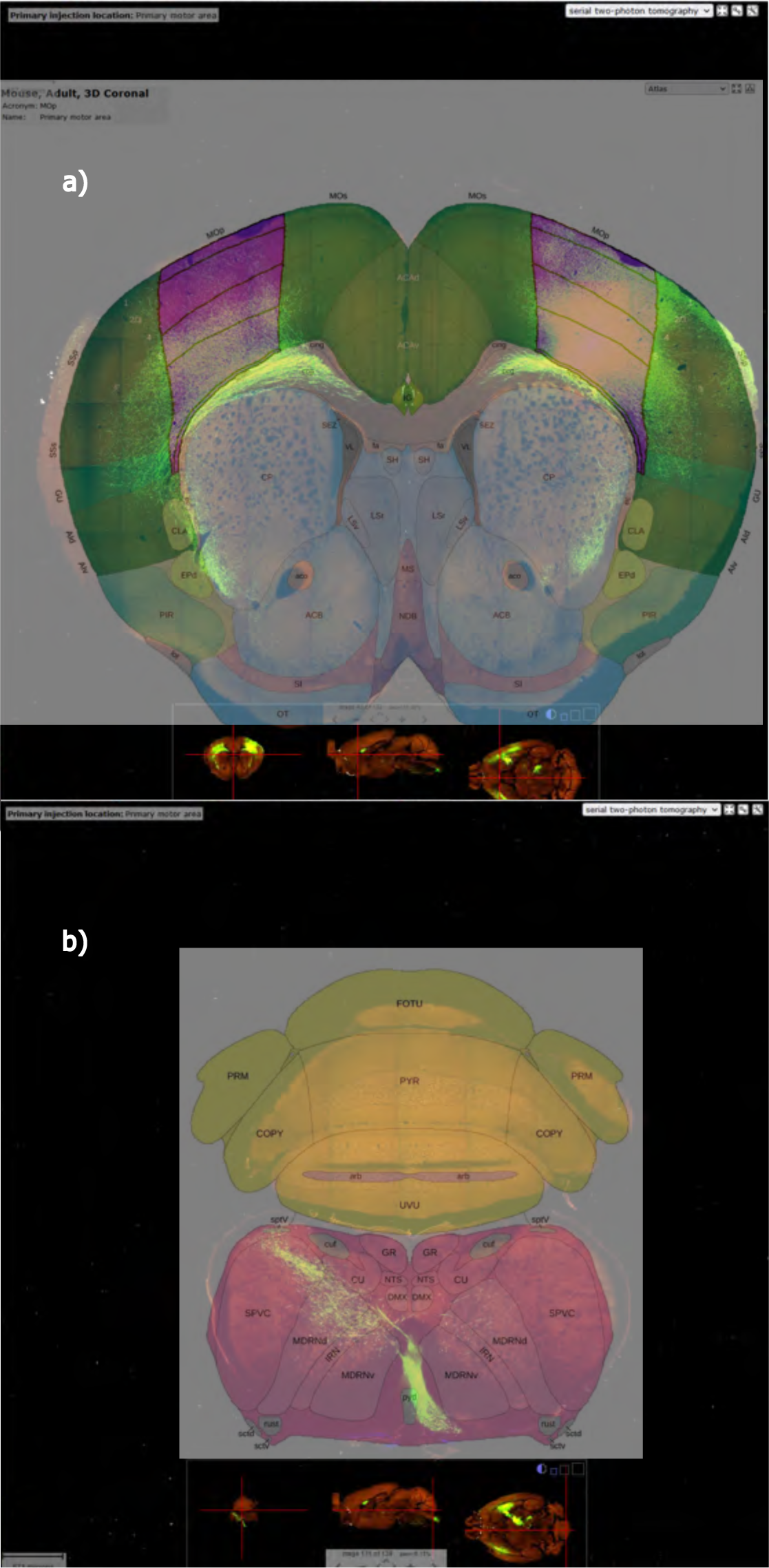
**a)** Injection structure MOp to **b)** SPVc. Experiment number 100141780, wild-type. The shortcomings of experimental procedures are addressed in the discussion.

**Supplemental Fig. 10.**
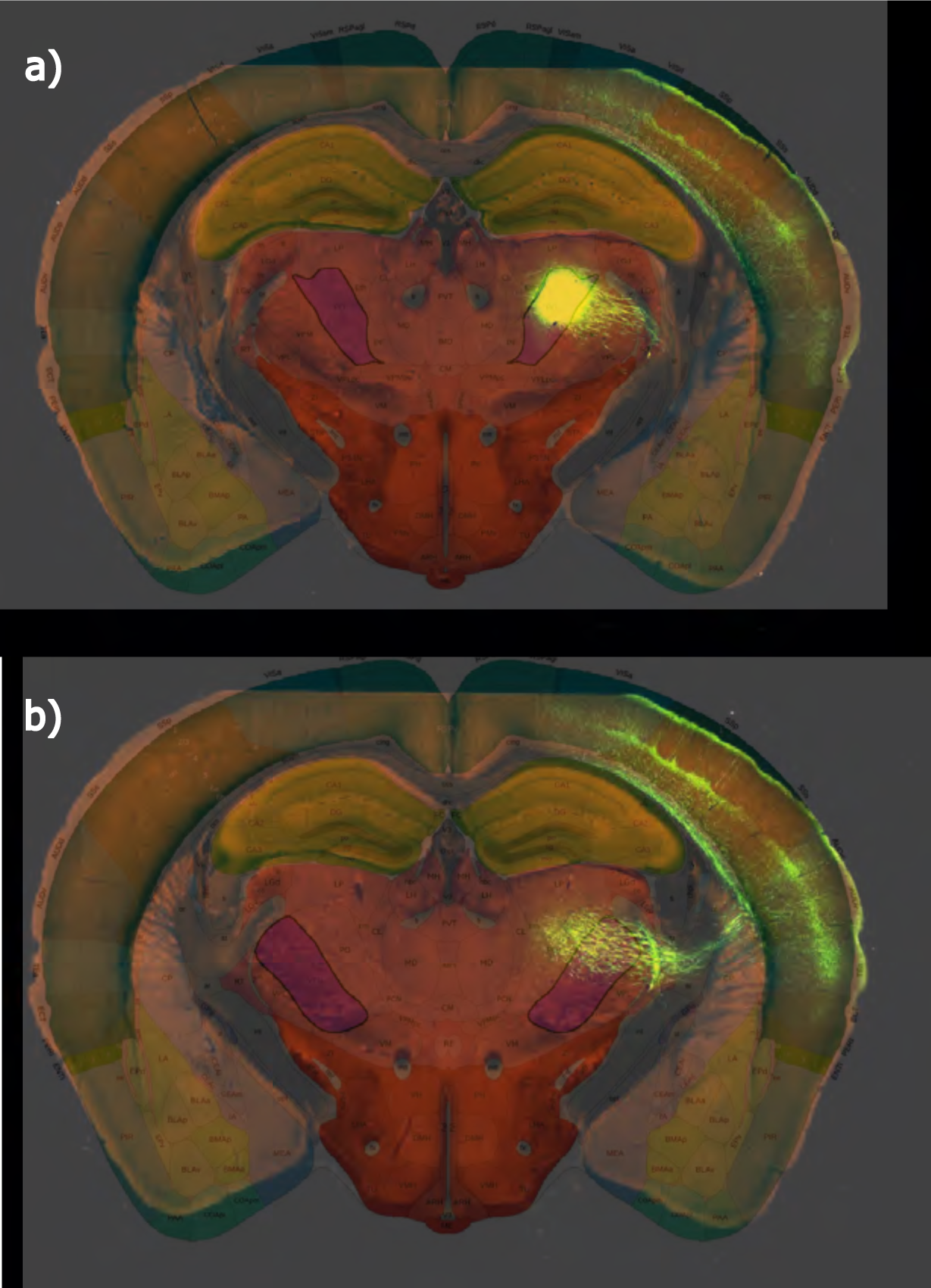
**a)** Injection structure PO to **b)** VPM. Experiment number 183011353, line Grik4-Cre. The shortcomings of experimental procedures are addressed in the discussion.

**Supplemental Fig. 11.**
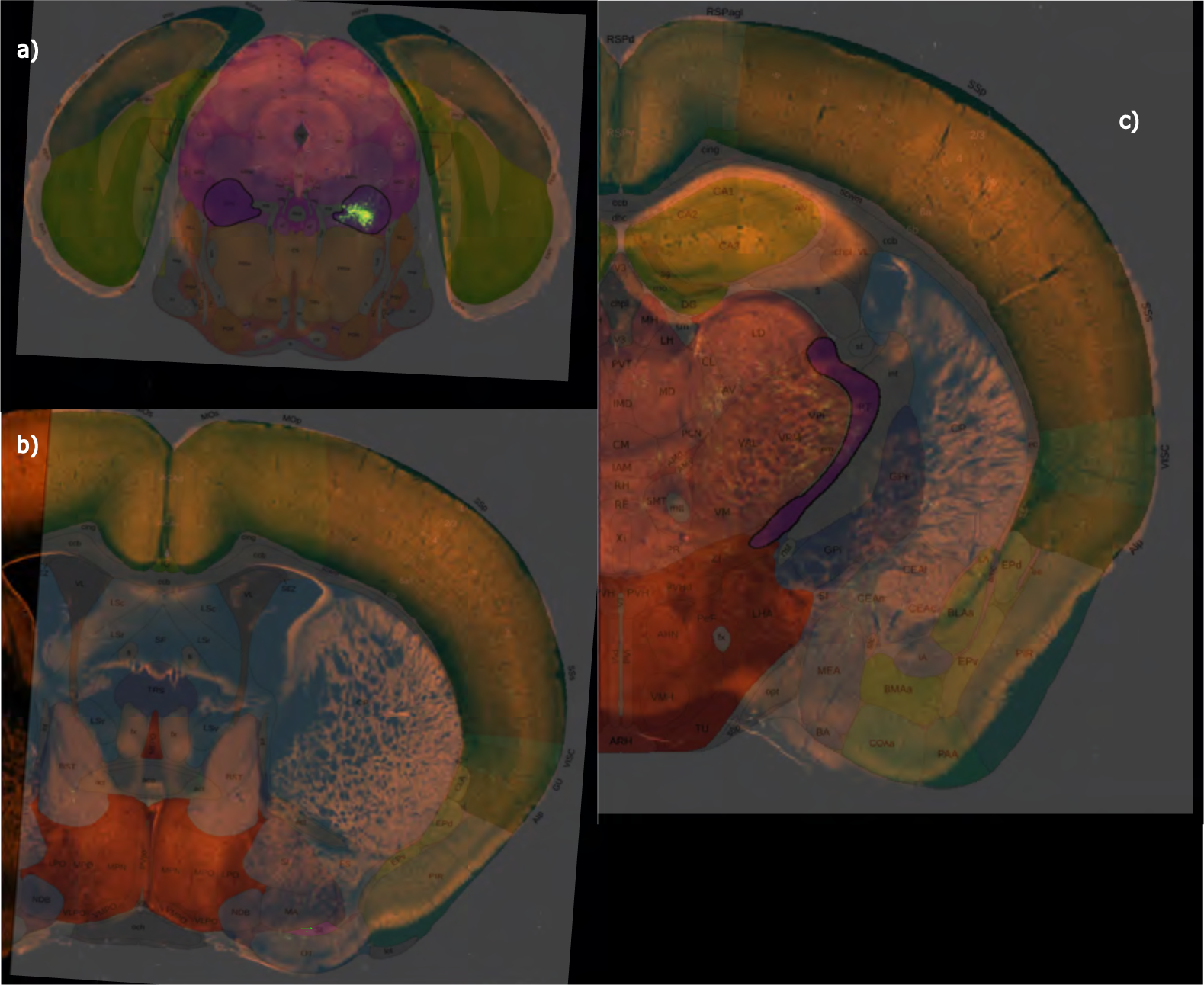
**a)** Injections structure PPN to **b)** SI and **c)** RT. Experiment number 264566672, line Chat-IRES-Cre-neo. The shortcomings of experimental procedures are addressed in the discussion.

**Supplemental Fig. 12.**
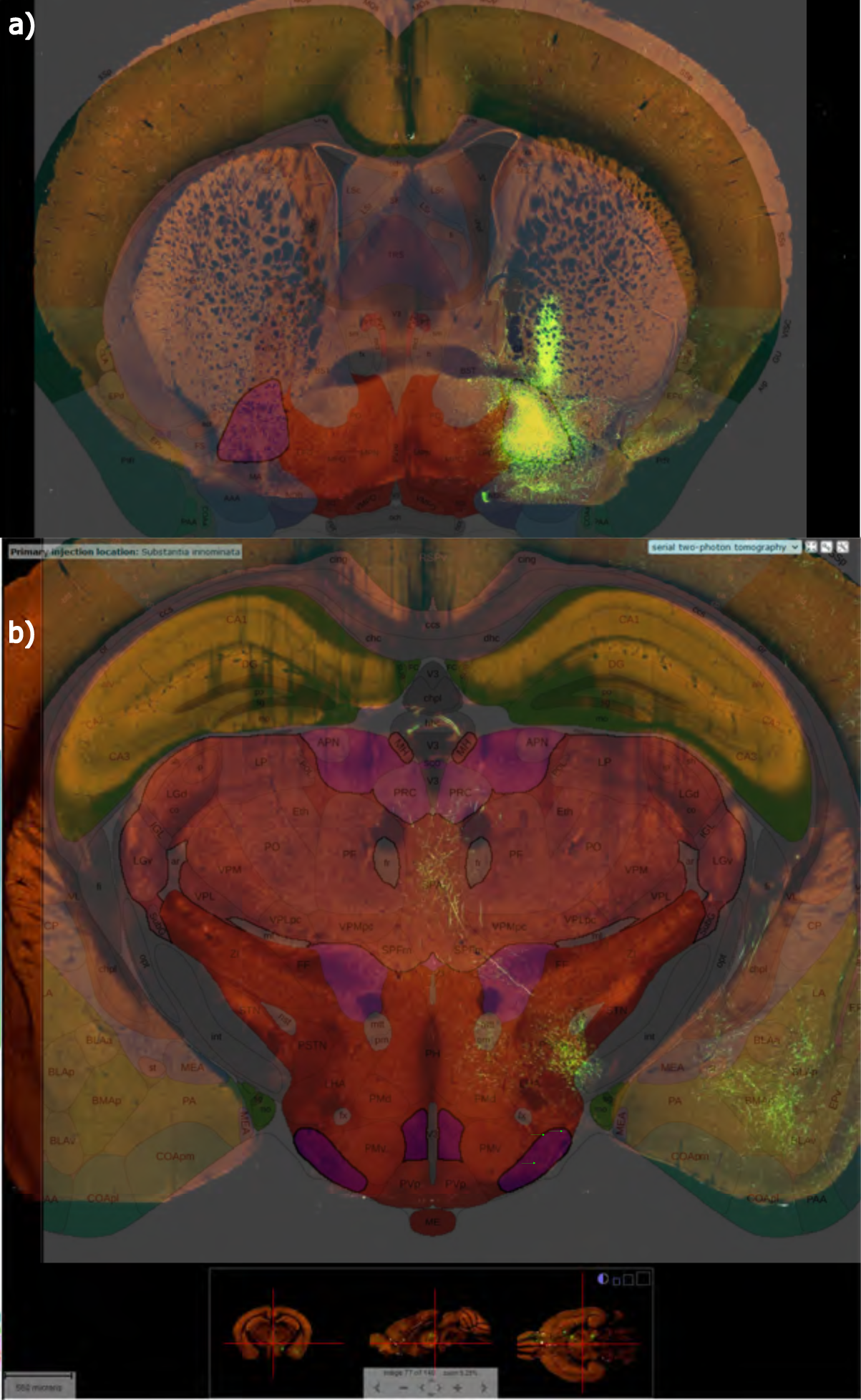
**a)** Injection structure SI to **b)** TM. Experiment number 305026861, line Drd3-Cre-KI196. The shortcomings of experimental procedures are addressed in the discussion.

**Supplemental Fig. 13.**
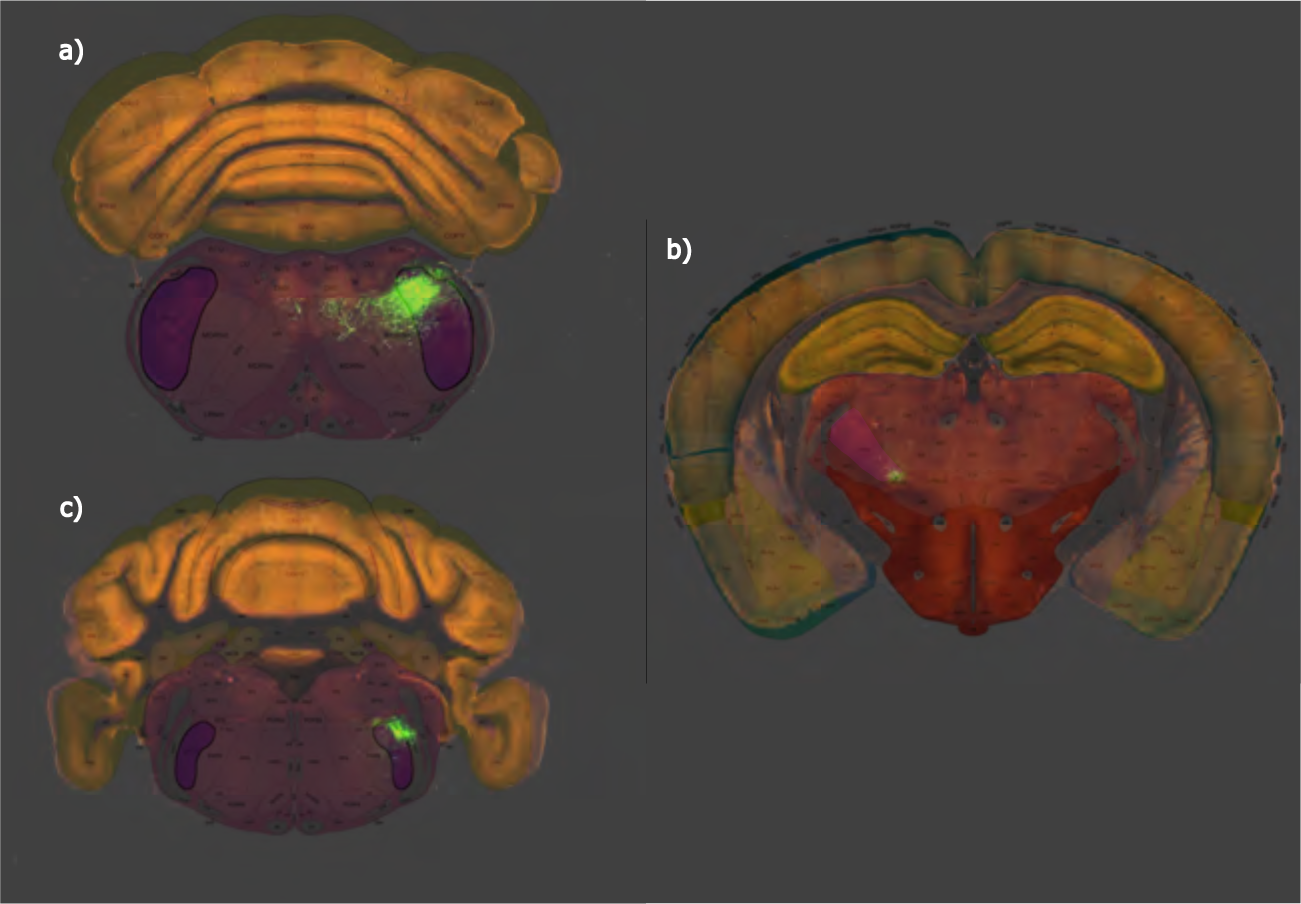
**a)** Injection structure SPVc to **b)** VPM and **c)** SPVo. Experiment number 114402050, wild-type. The shortcomings of experimental procedures are addressed in the discussion.

**Supplemental Fig. 14.**
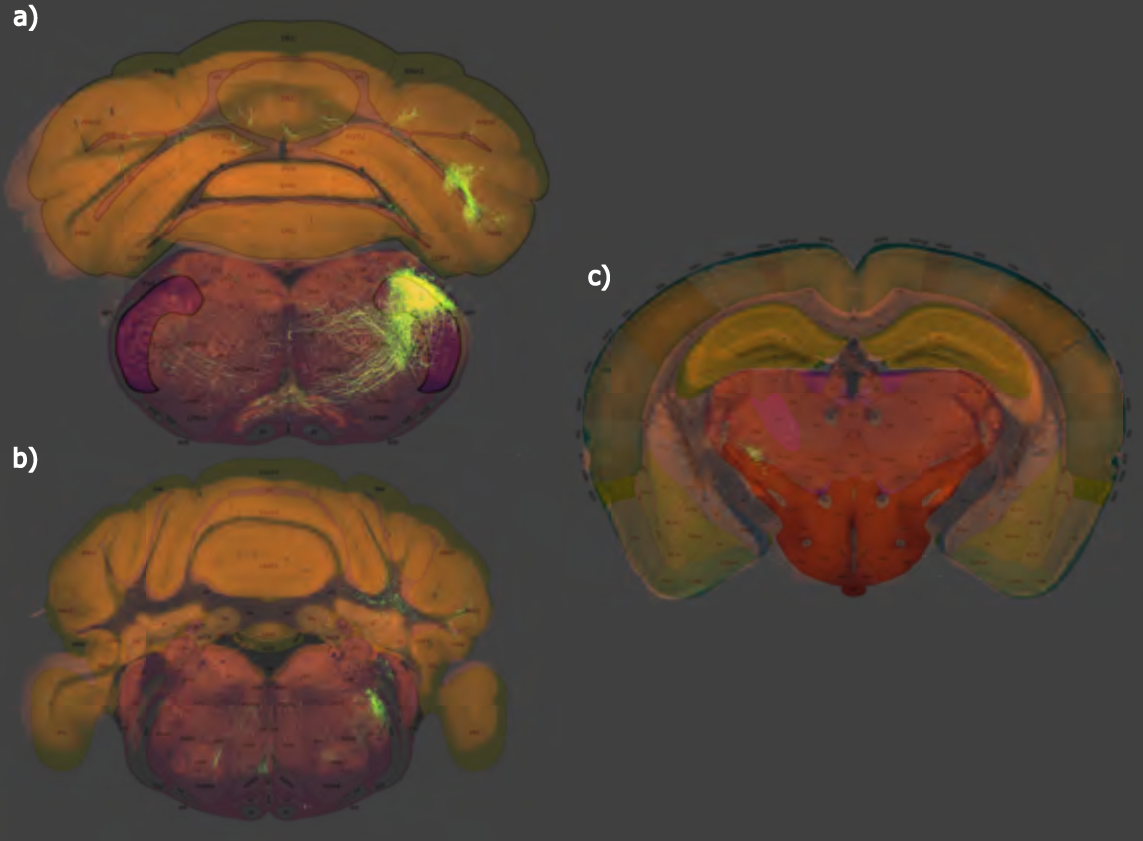
**a)** Injection structure SPVi to **b)** SPVo and **c)** PO. Experiment number 272738620, wild-type. The shortcomings of experimental procedures are addressed in the discussion.

**Supplemental Fig. 15.**
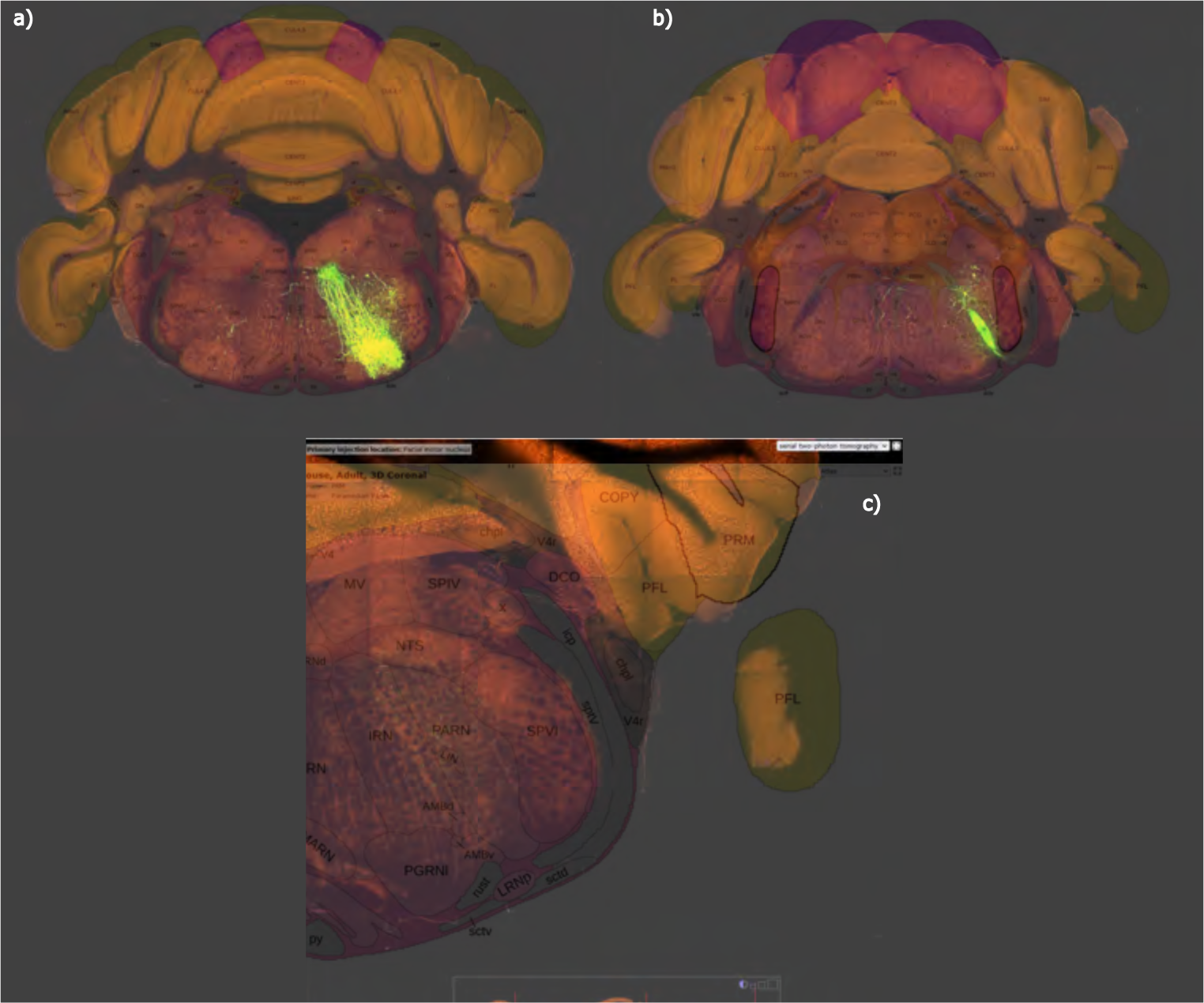
**a)** Injection structure VII to SPVo **b)** PSV, and **c)** to SPVi. Experiment number 177905562, line Scnn1a-Tg2-Cre. The shortcomings of experimental procedures are addressed in the discussion.

**Supplemental Fig. 16.**
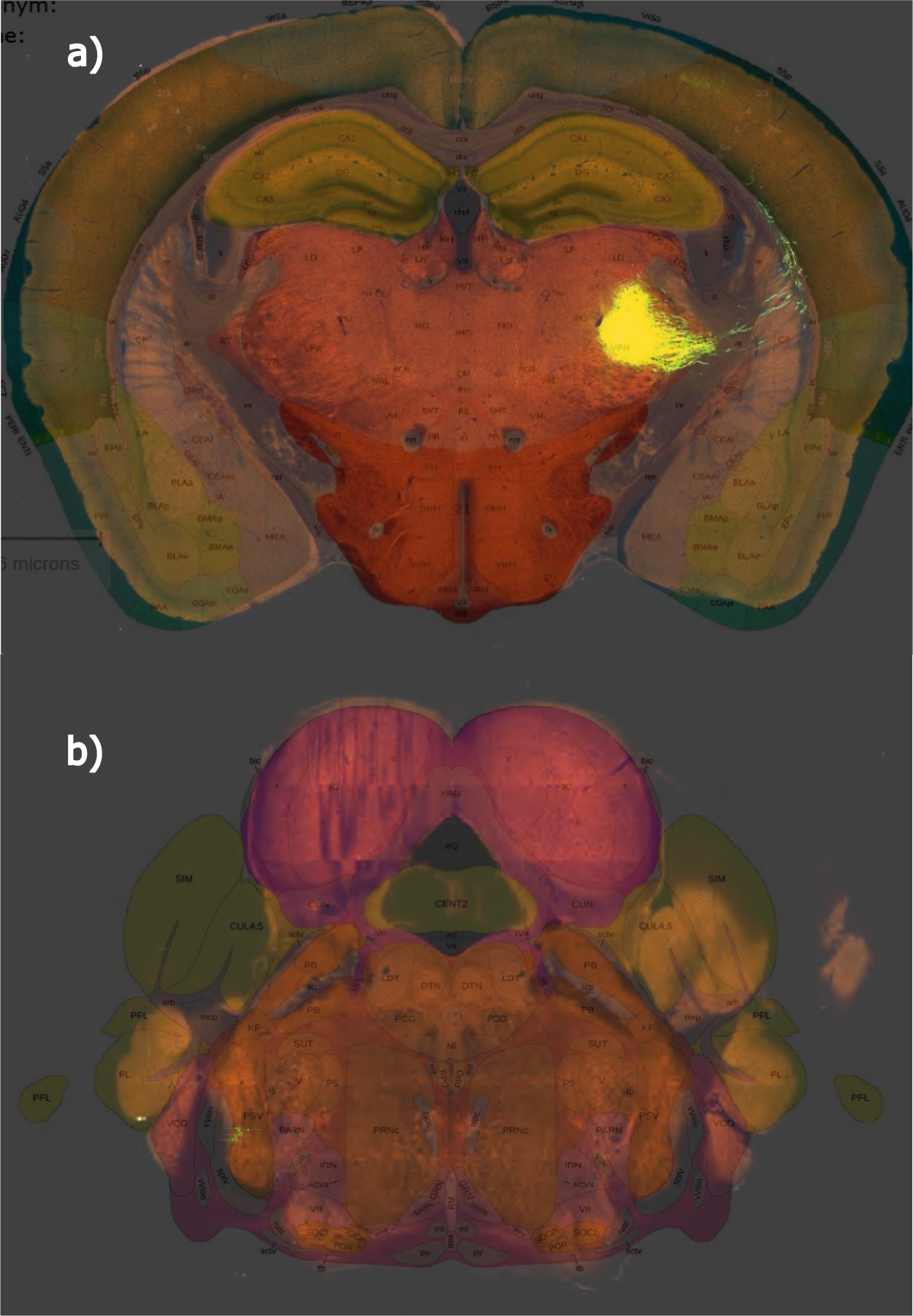
**a)** Injection structure VPM to **b)** PSV, experiment number 312240825, line Slc17a6-IRES-Cre. The shortcomings of experimental procedures are addressed in the discussion.

**Supplemental Fig. 17.**
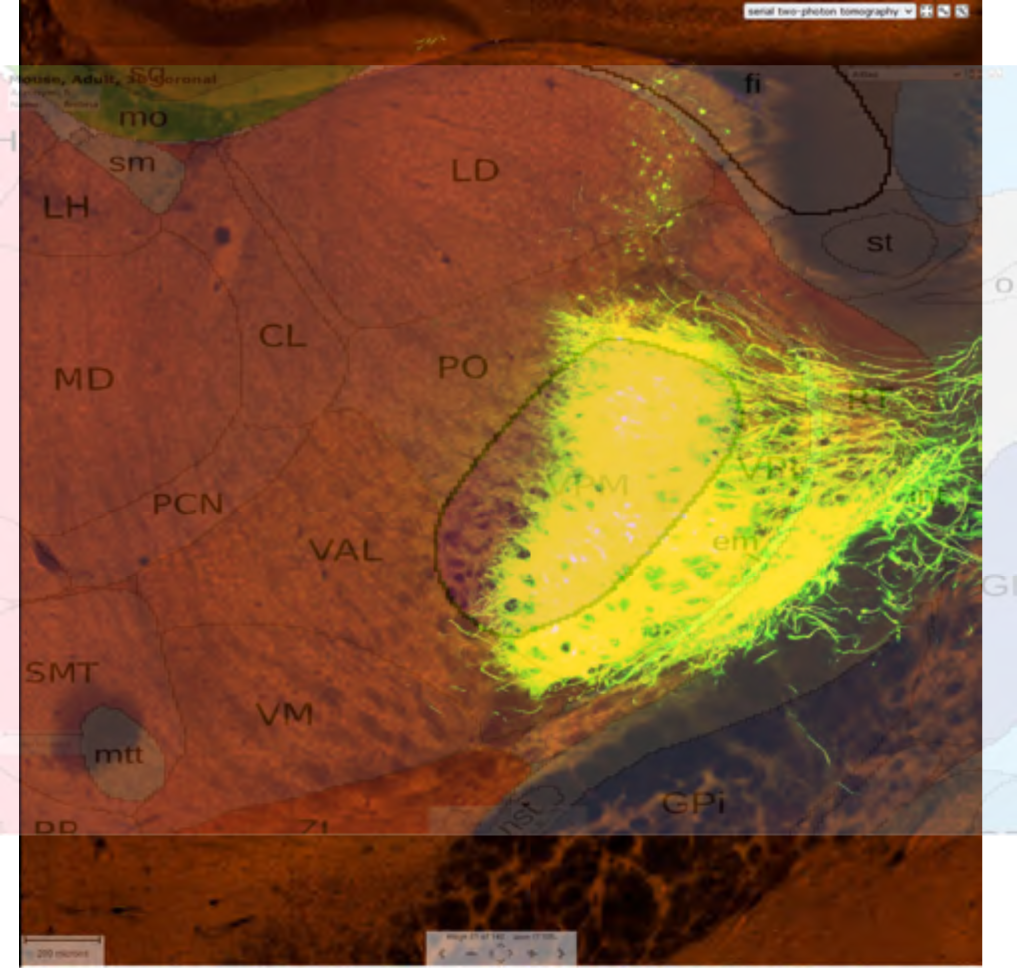
Inhibition target and source VPM to PO, experiment number 268206050, line Ppp1r17-Cre-NL146. The shortcomings of experimental procedures are addressed in the discussion.

**Supplemental Fig. 18.**
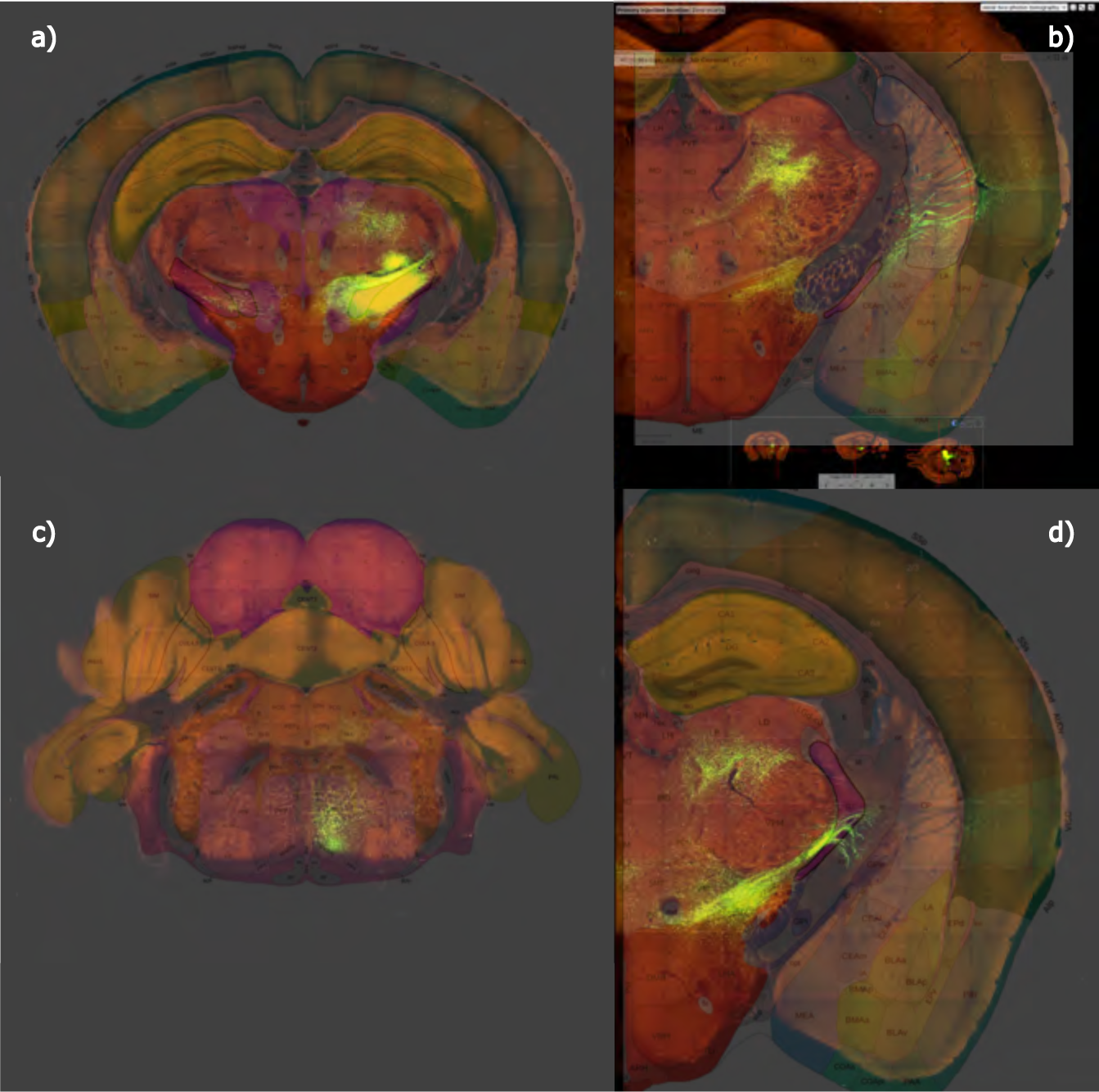
**a)** Injection structure ZI to **b)** SI, **c)** PSV, **d)** RT. Experiment number 113095845, wild-type. The shortcomings of experimental procedures are addressed in the discussion.

**Supplemental Fig. 19.**
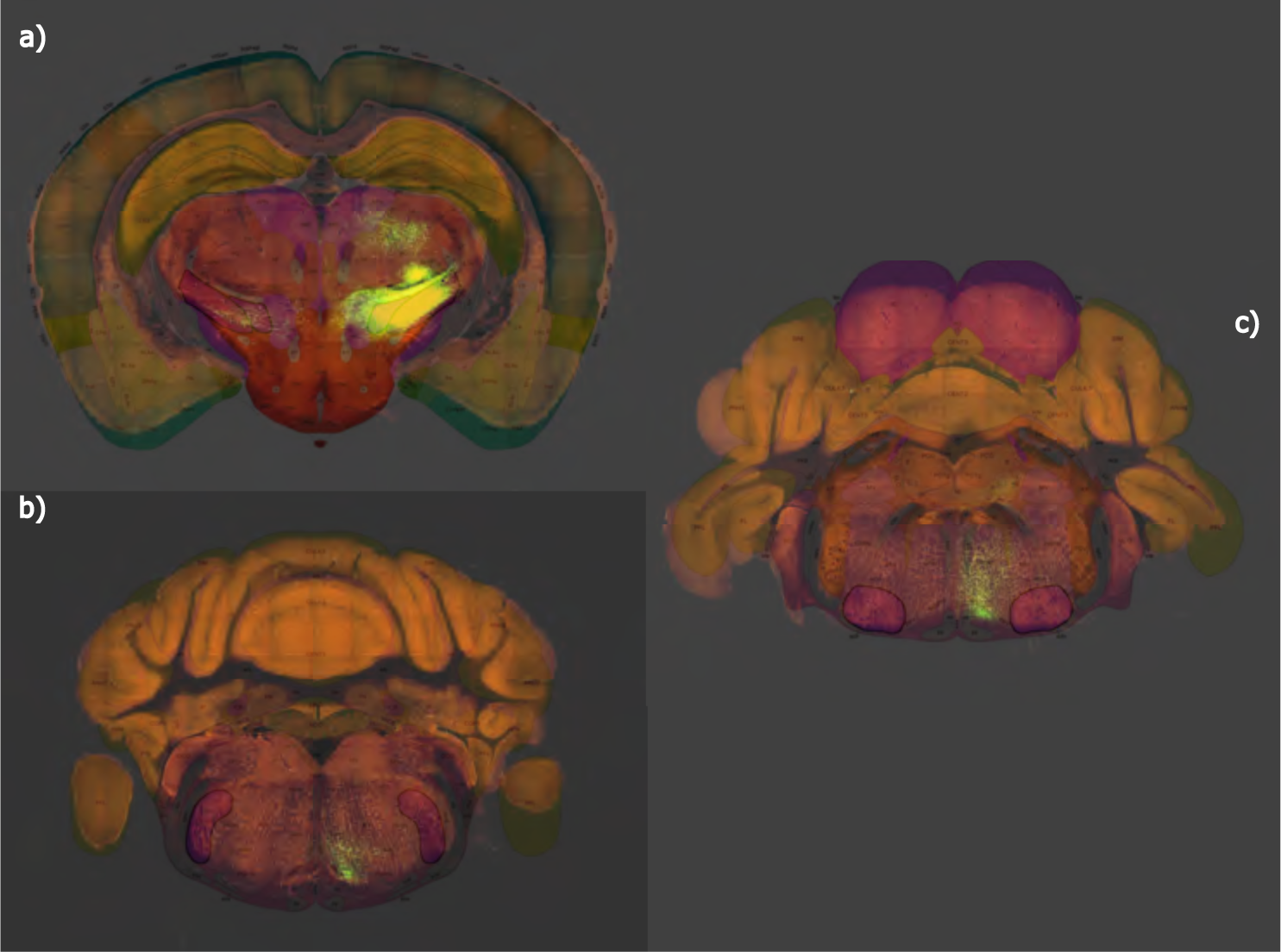
**a)** Injection structure ZI to **b)** SPVo, and **c)** VII. Experiment number 113095845, wild-type. The shortcomings of experimental procedures are addressed in the discussion.

**Supplemental Fig. 20.**
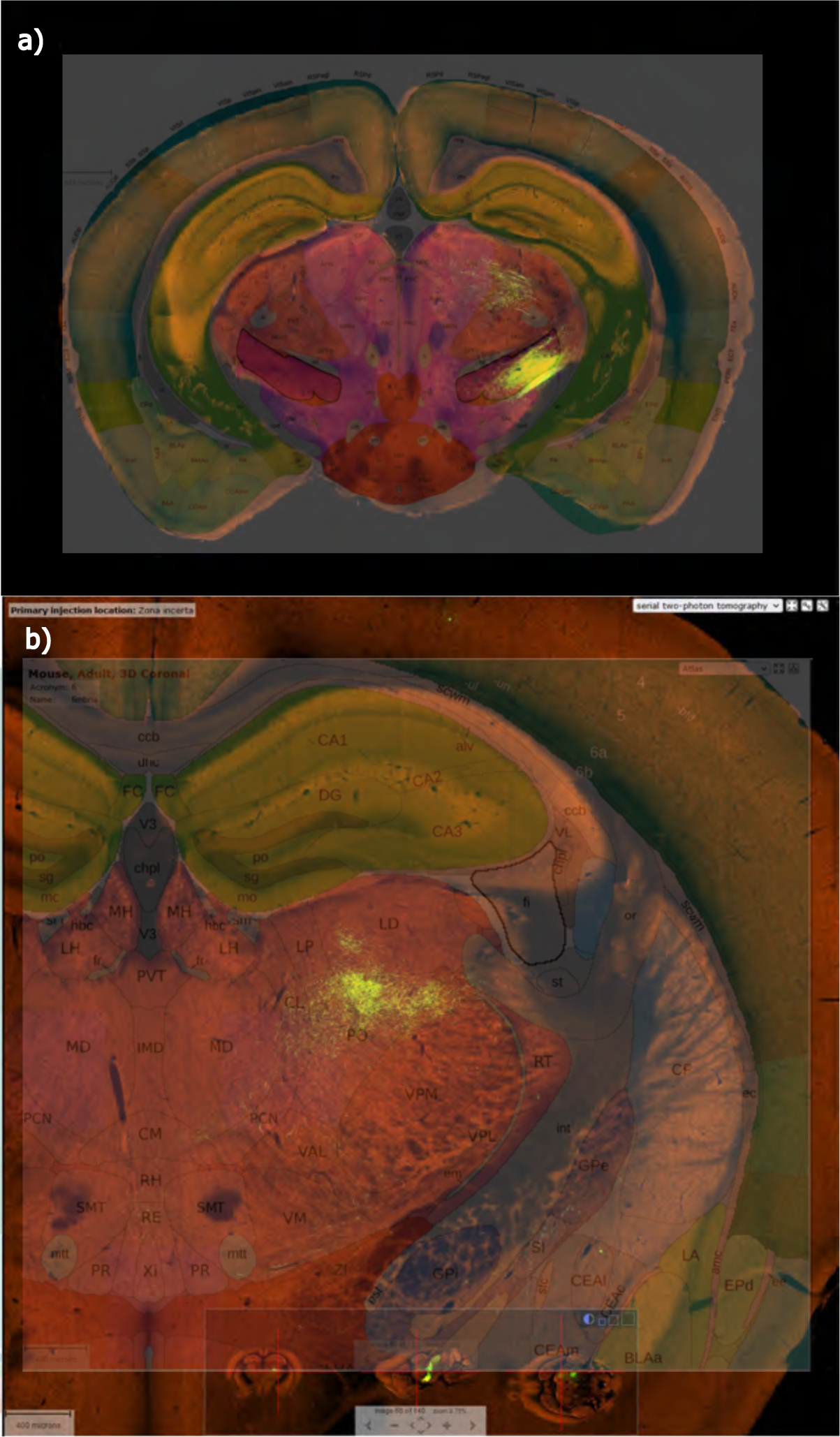
**a)** Injection structure ZI to **b)** VPM. Experiment number 301539438, line Pvalb-IRES-Cre. The shortcomings of experimental procedures are addressed in the discussion.

**Supplemental Fig. 21.**
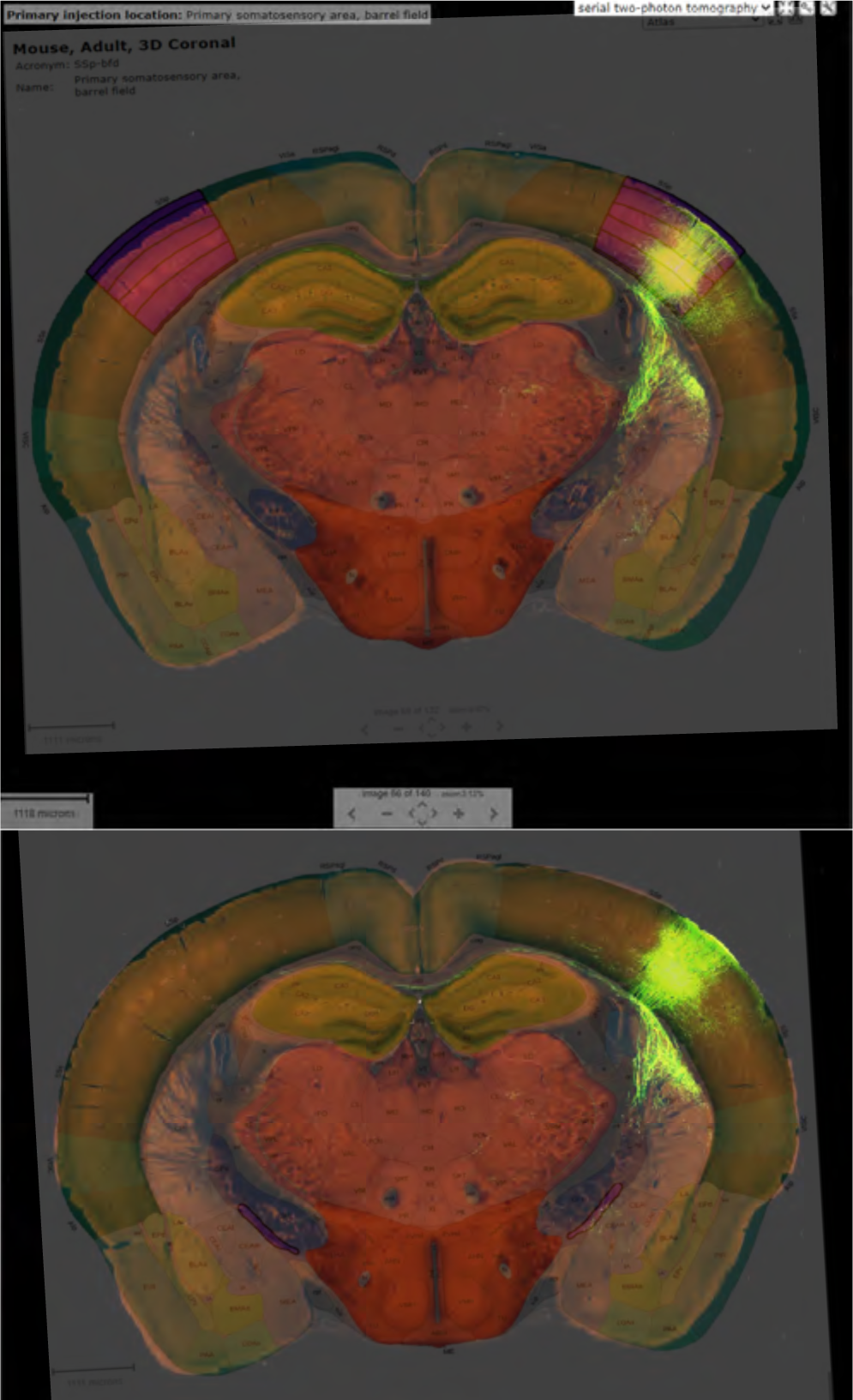
**a)** Injection structure bfd to **b)** SI. Experiment number 159602992, line Cart-Tg1-Cre. The shortcomings of experimental procedures are addressed in the discussion.

**Supplemental Fig. 22.**
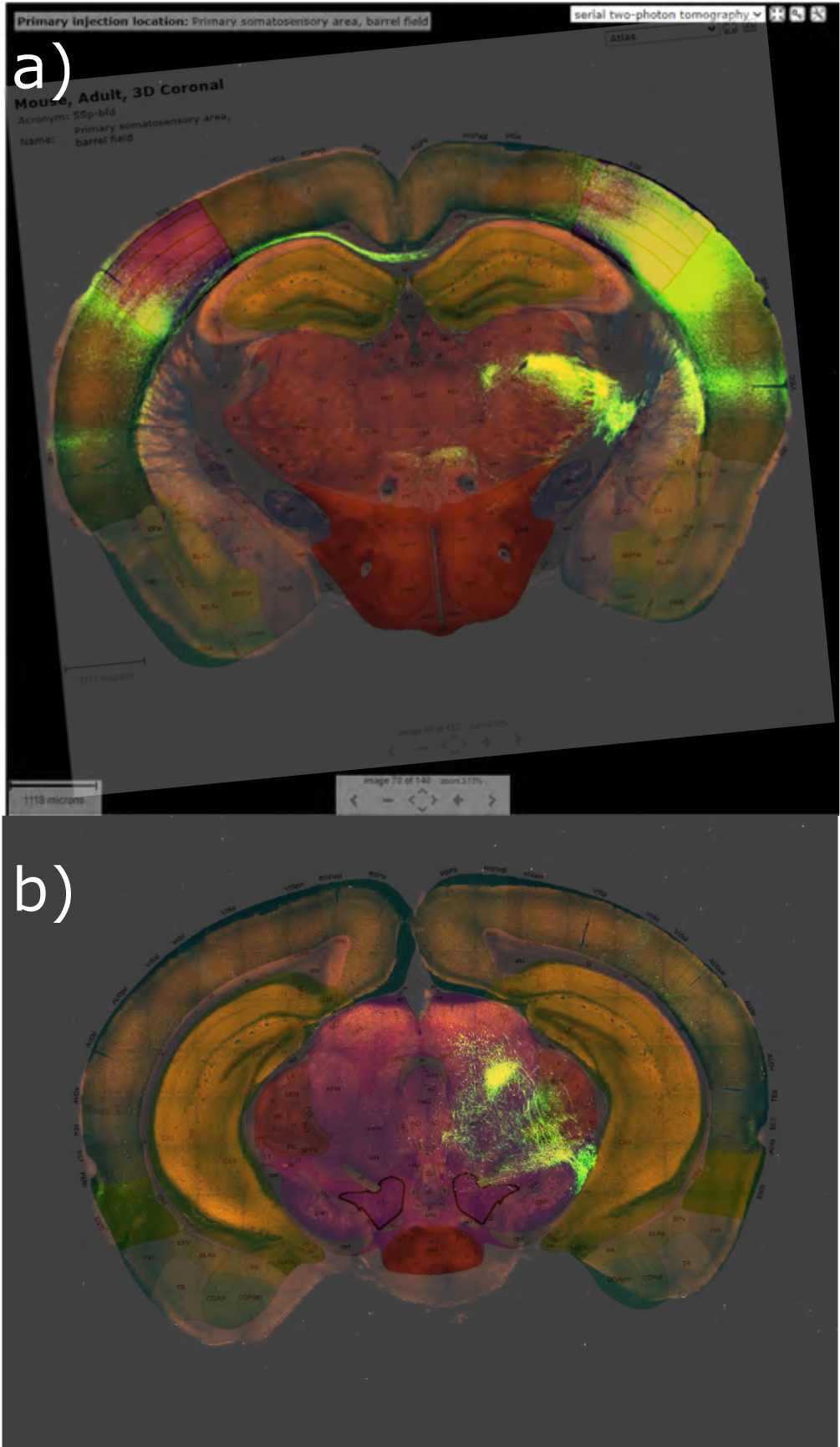
**a)** Injection structure bfd to **b)** VTA. Experiment number 112951804, wild-type. The shortcomings of experimental procedures are addressed in the discussion.

**Supplemental Fig. 23.**
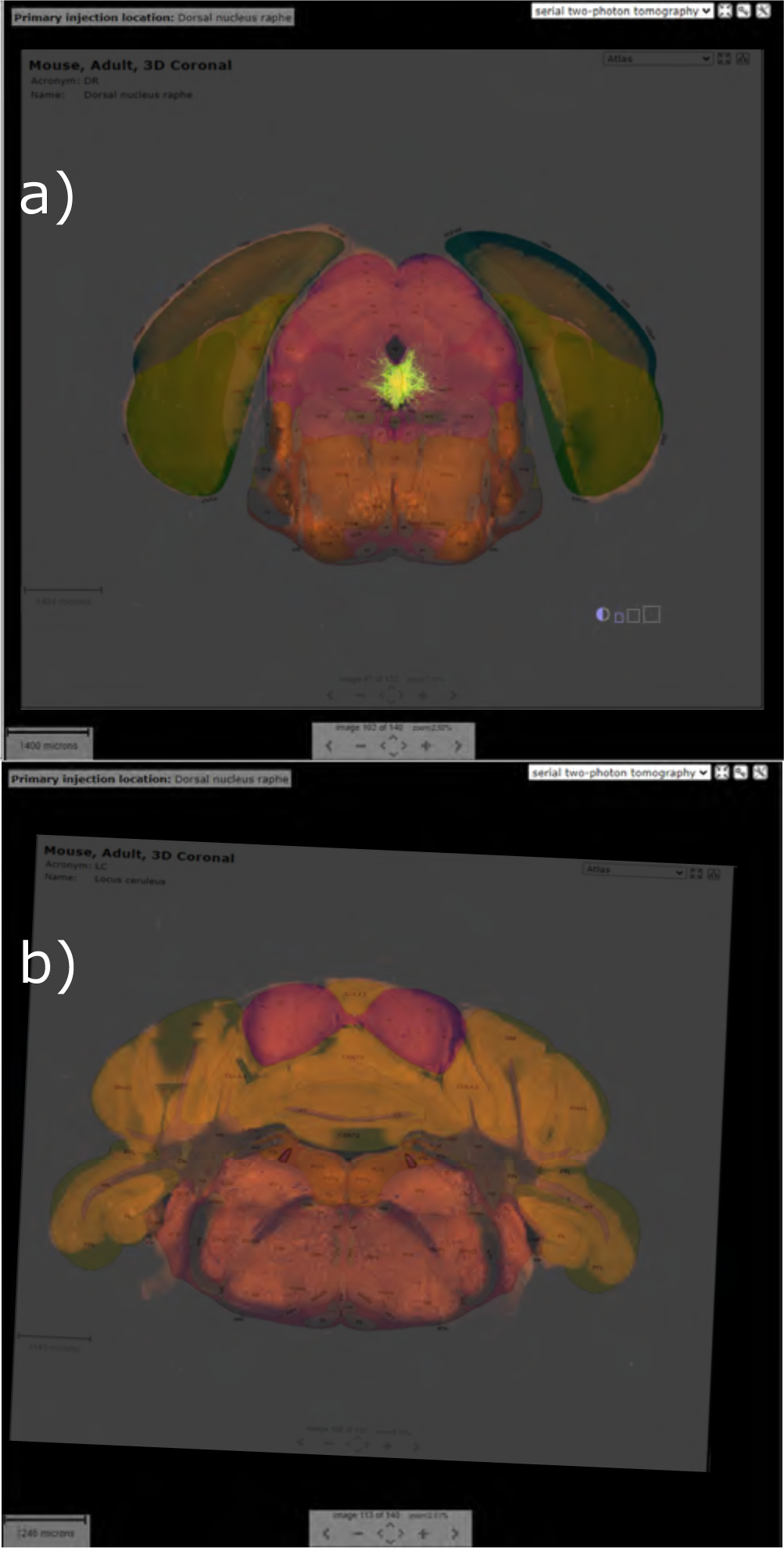
**a)** Injection structure DR to **b)** LC. Experiment number 547510030, Slc17a8-IRES2-Cre. The shortcomings of experimental procedures are addressed in the discussion.

**Supplemental Fig. 24.**
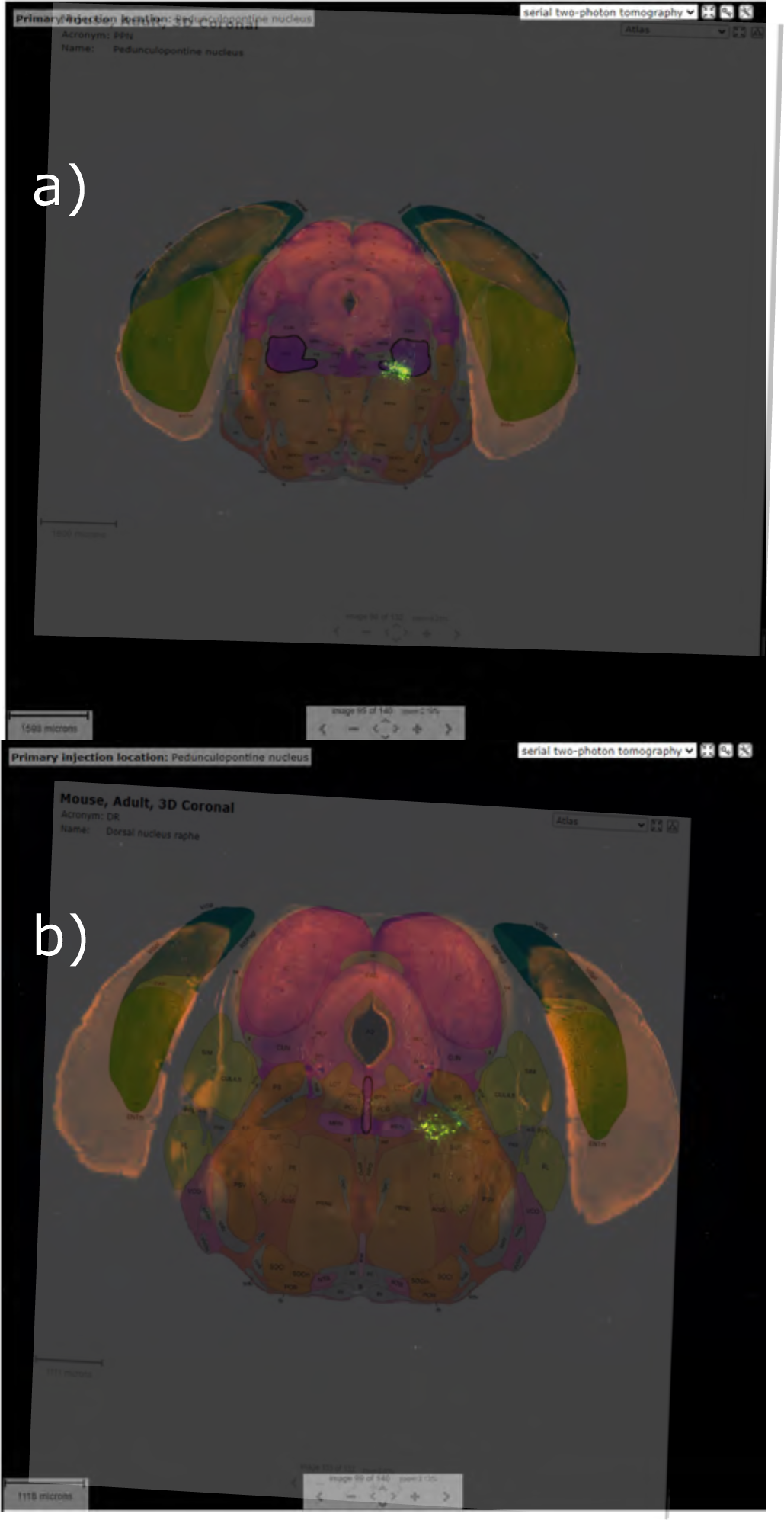
**a)** Injection structure PPN to **b)** DR. Experiment number 264566672, Chat-IRES-Cre-neo. The shortcomings of experimental procedures are addressed in the discussion.

**Supplemental Fig. 25.**
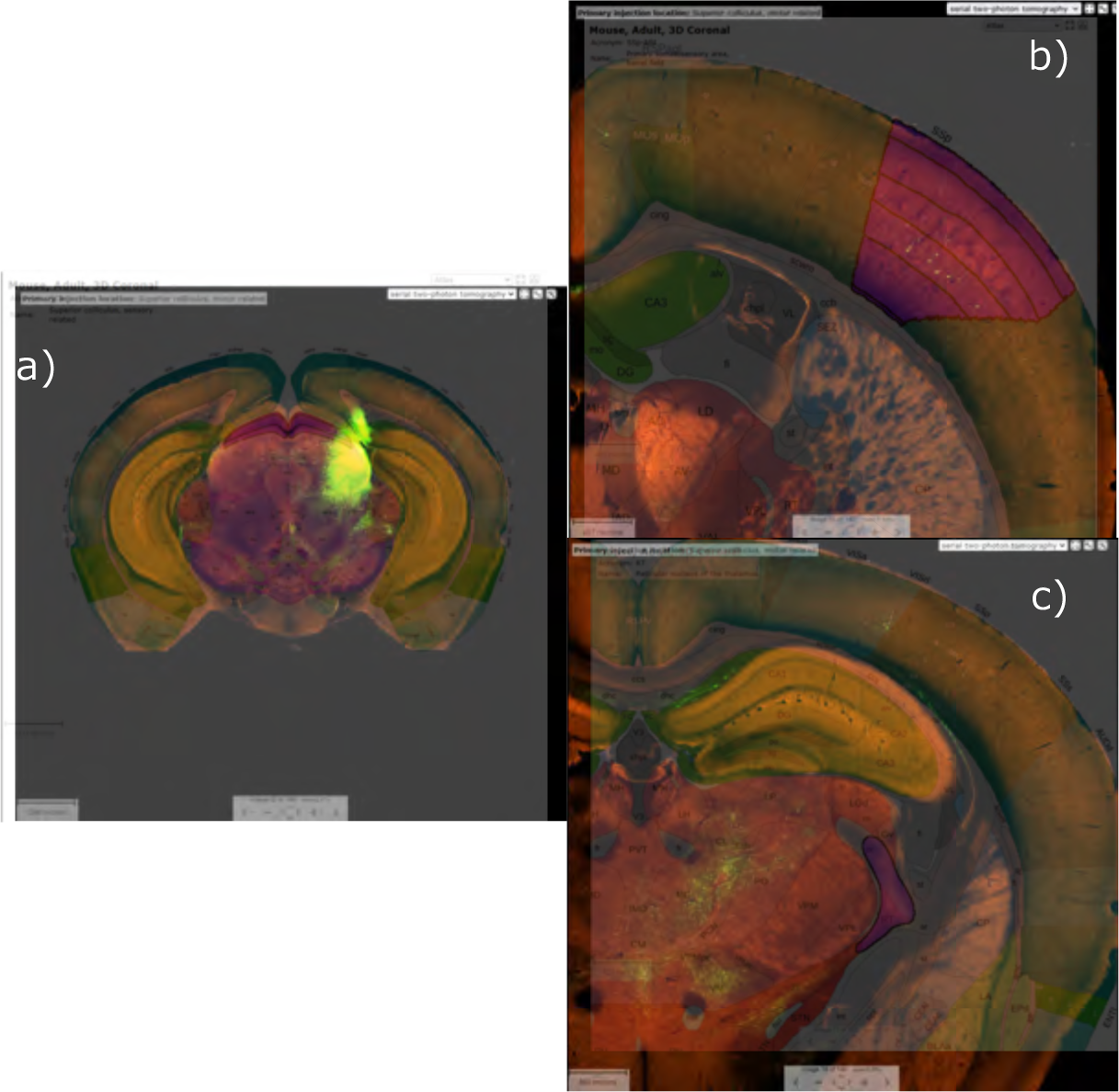
**a)** Injection structure SC to **b)** bfd and **c)** DR. Experiment number 146078721, wild-type. The shortcomings of experimental procedures are addressed in the discussion.

**Supplemental Fig. 26.**
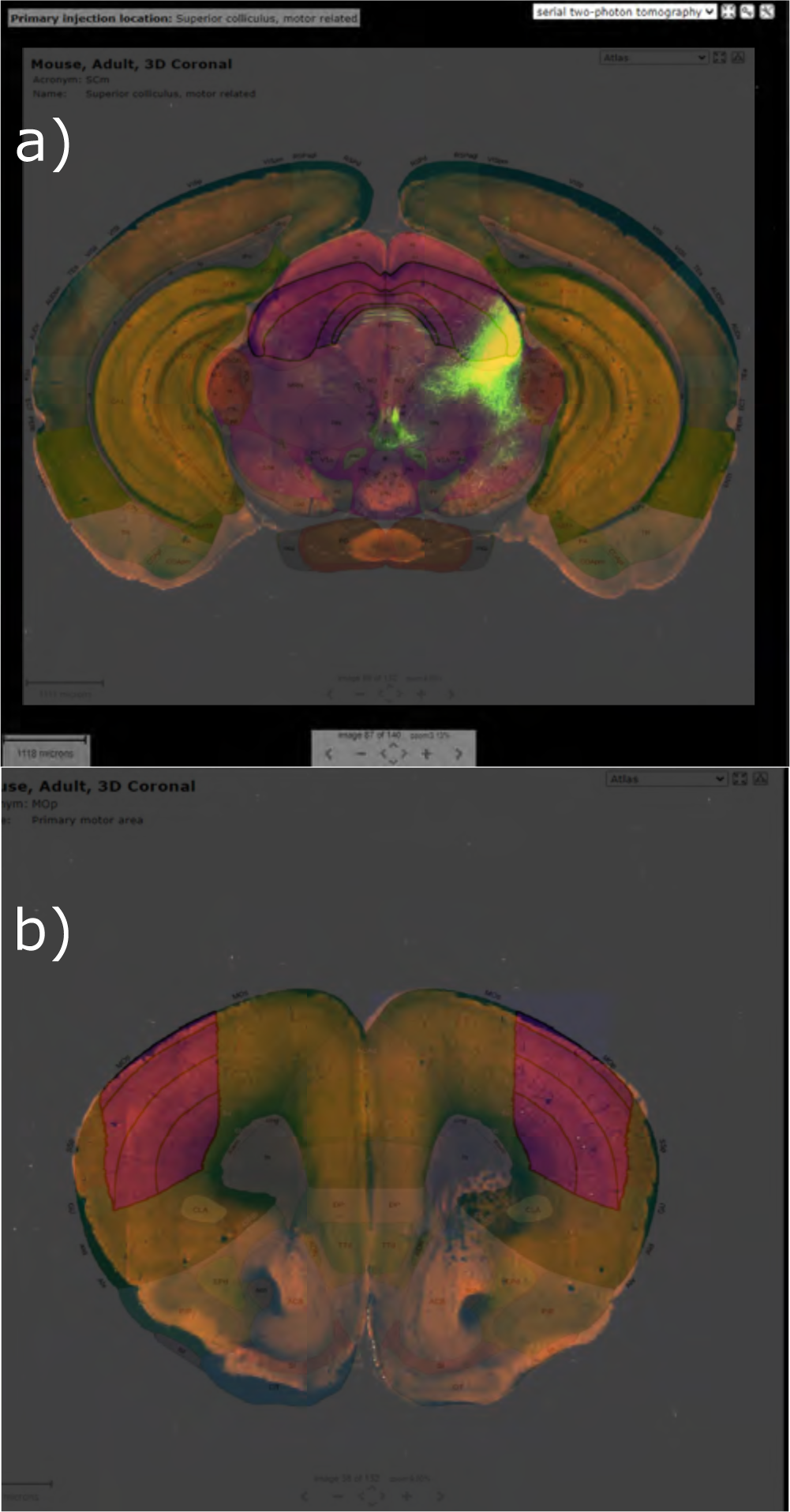
**a)** Injection structure SC to **b)** MOp. Experiment number 175158132, wild-type. The shortcomings of experimental procedures are addressed in the discussion.

**Supplemental Fig. 27.**
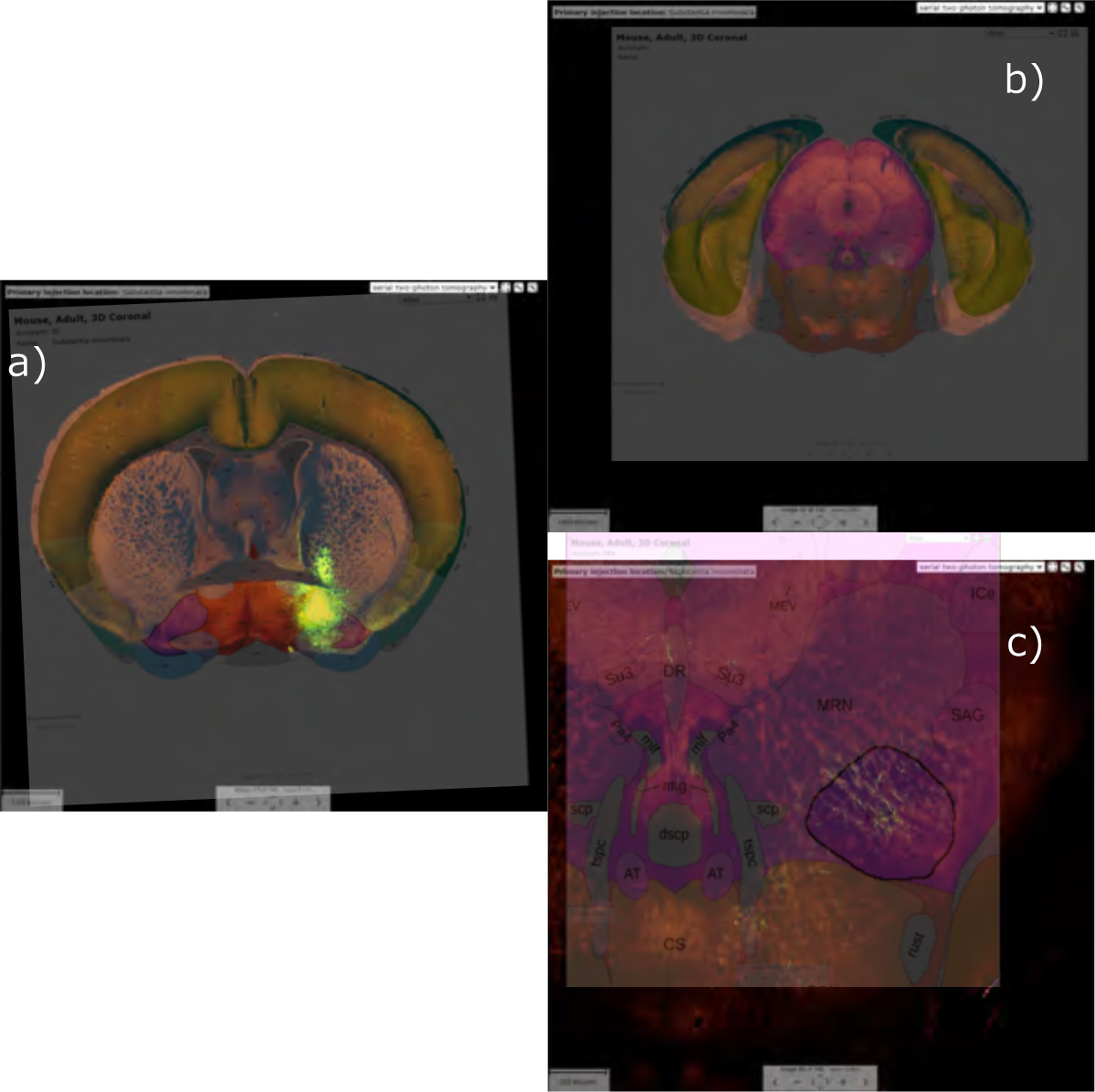
**a)** Injection structure SI to **b)** PPN. **c)** zoom on PPN. Experiment number 305026861, Drd3-Cre-KI196. The shortcomings of experimental procedures are addressed in the discussion.

**Supplemental Fig. 28.**
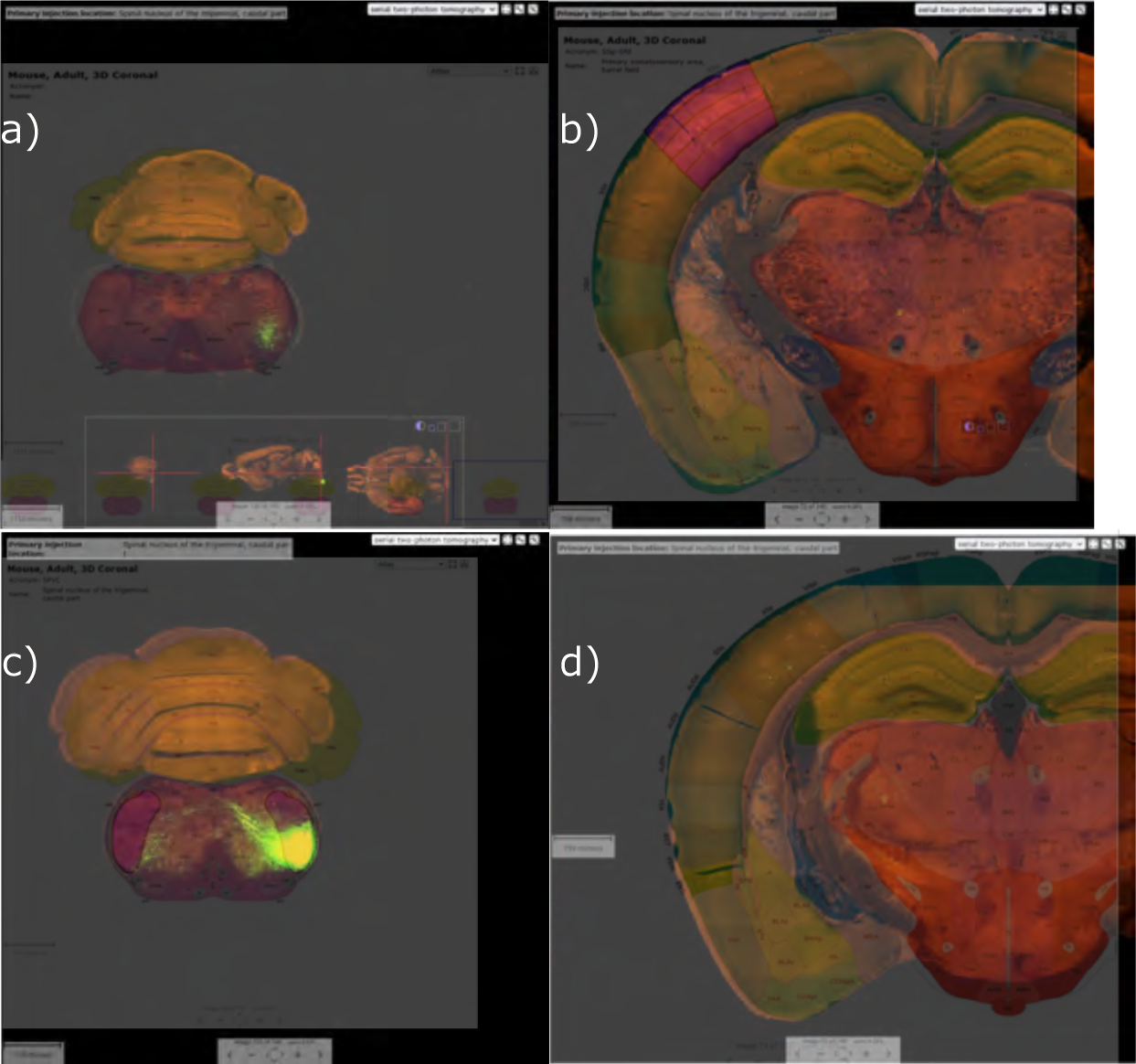
**a)** Injection structure SPVc to **b)** bfd. **c)** Injection structure SPVc to **d)** bfd. a) and b) experiment number 287460307, Cck-IRES-Cre. c) and d) experiment number 147215105, wild-type. The shortcomings of experimental procedures are addressed in the discussion.

**Supplemental Fig. 29.**
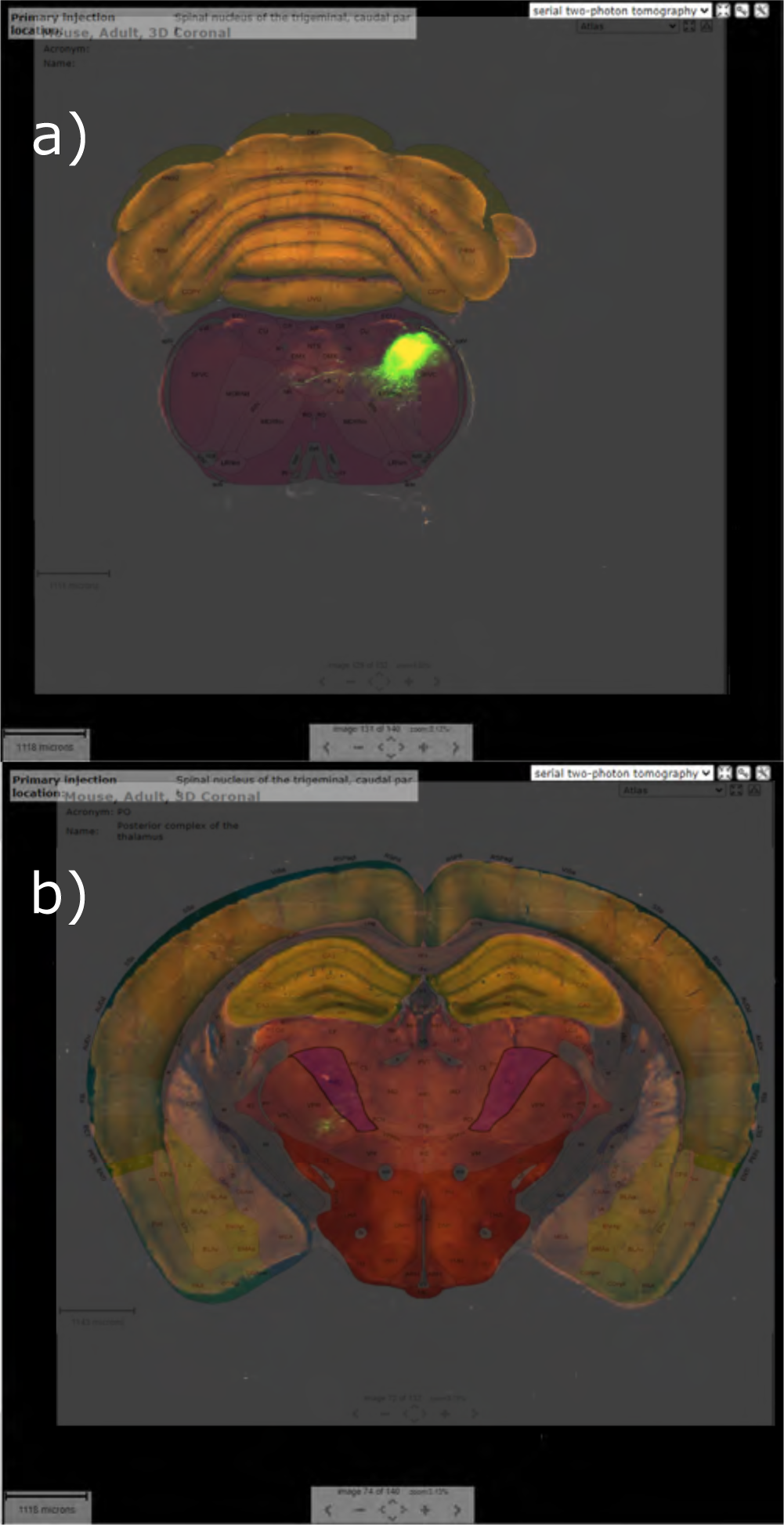
**a)** Injection structure SPVc to **b)** PO. Experiment number 114402050, wild-type. The shortcomings of experimental procedures are addressed in the discussion.

**Supplemental Fig. 30.**
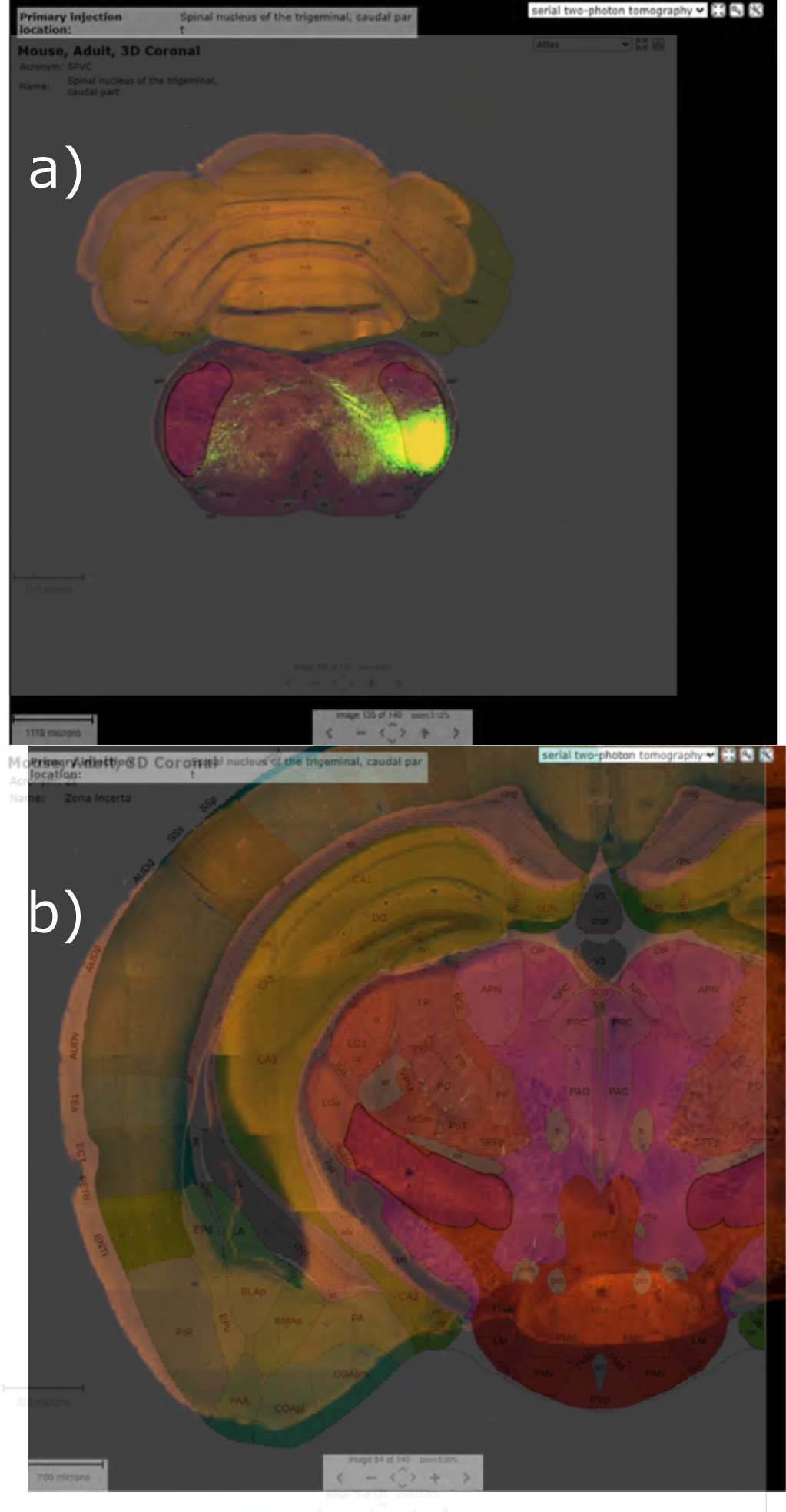
**a)** Injection structure SPVc to **b)** ZI. Experiment number 147215105, wild-type. The shortcomings of experimental procedures are addressed in the discussion.

**Supplemental Fig. 31.**
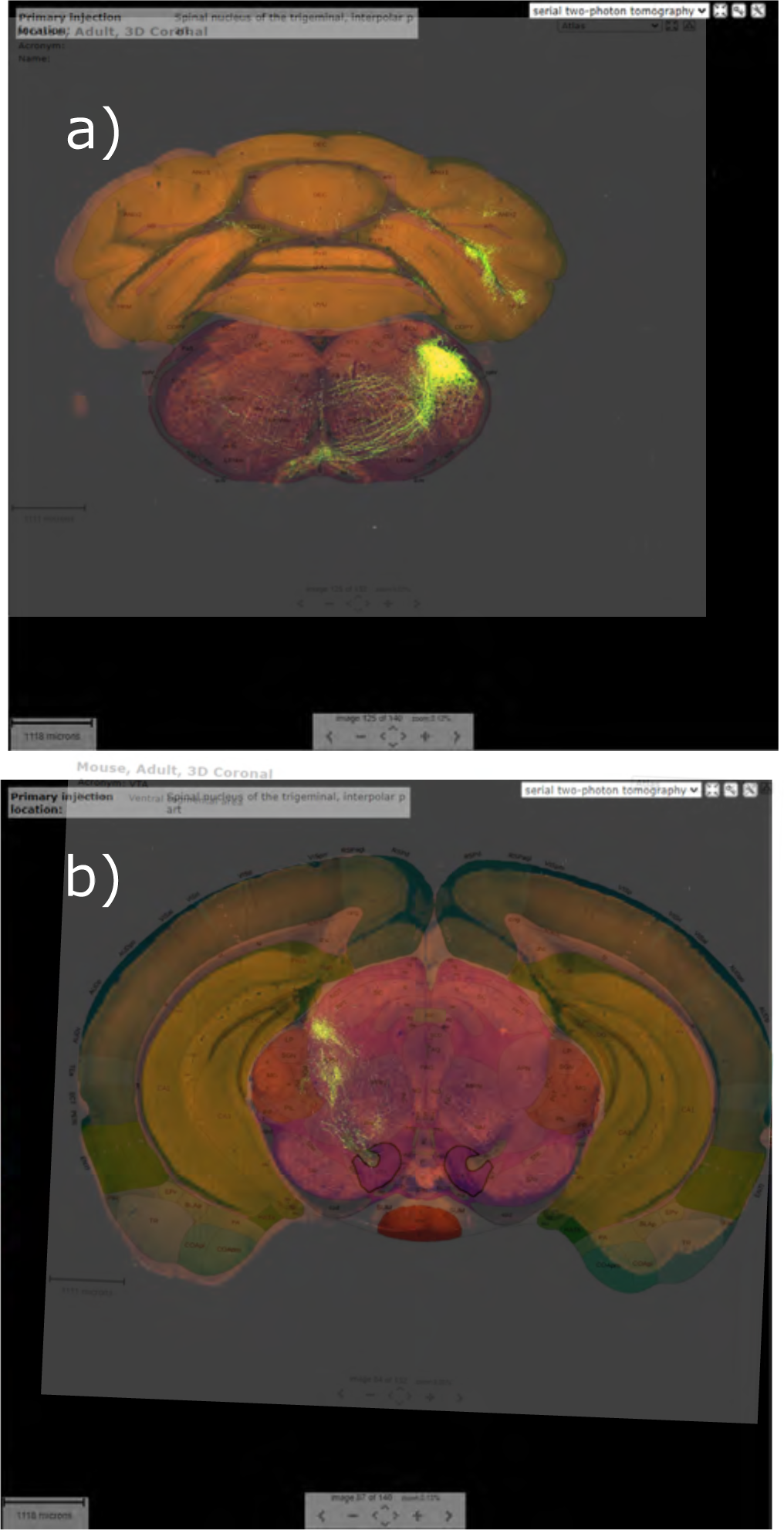
**a)** Injection structure SPVi to **b)** VTA. Experiment number 272738620, wild-type. The shortcomings of experimental procedures are addressed in the discussion.

**Supplemental Fig. 32.**
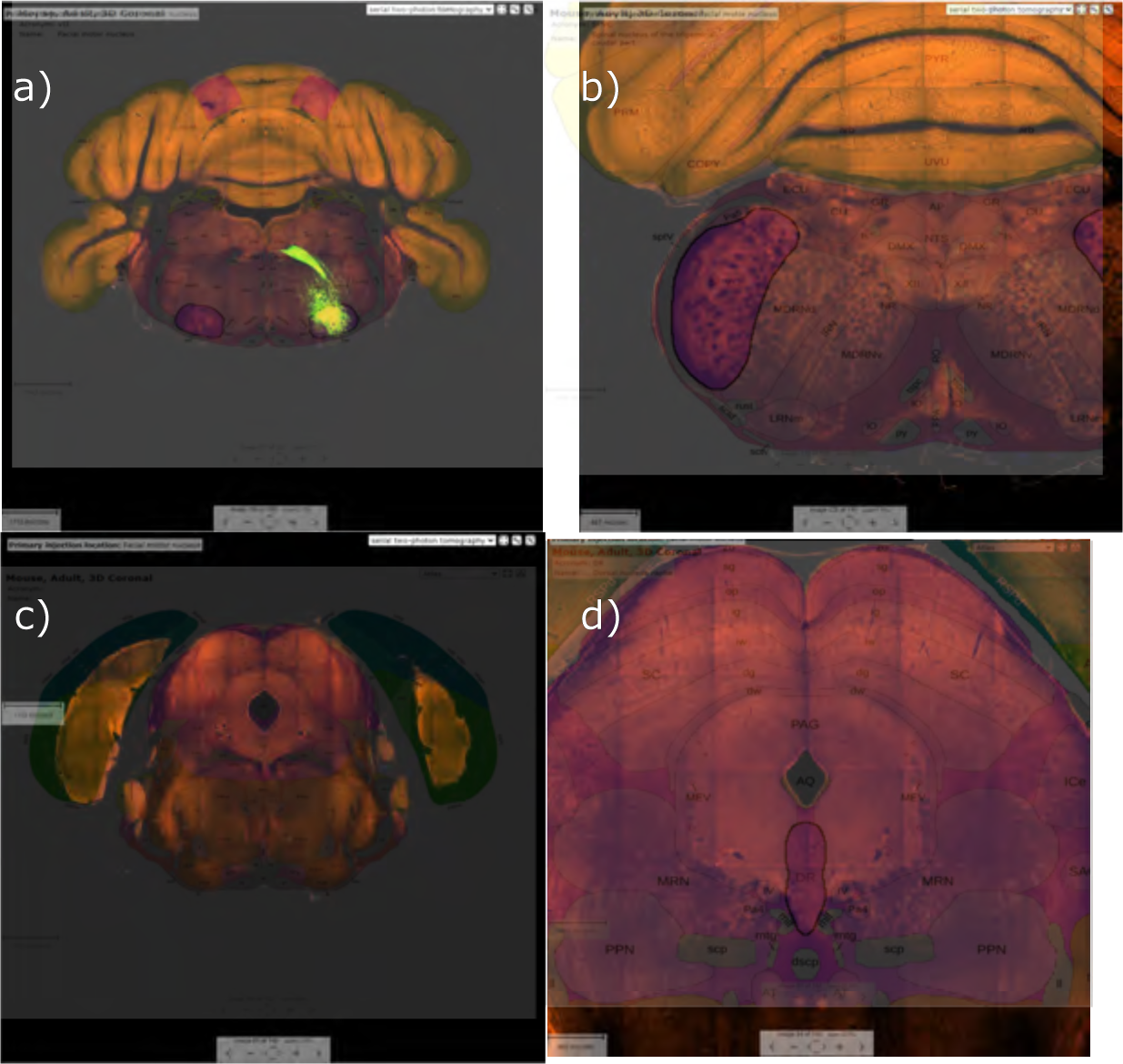
**a)** Injection structure VII to **b)** SPVc, **c)** SC and **d)** DR. Experiment number 113766744, Grik4-Cre. The shortcomings of experimental procedures are addressed in the discussion.

**Supplemental Fig. 33.**
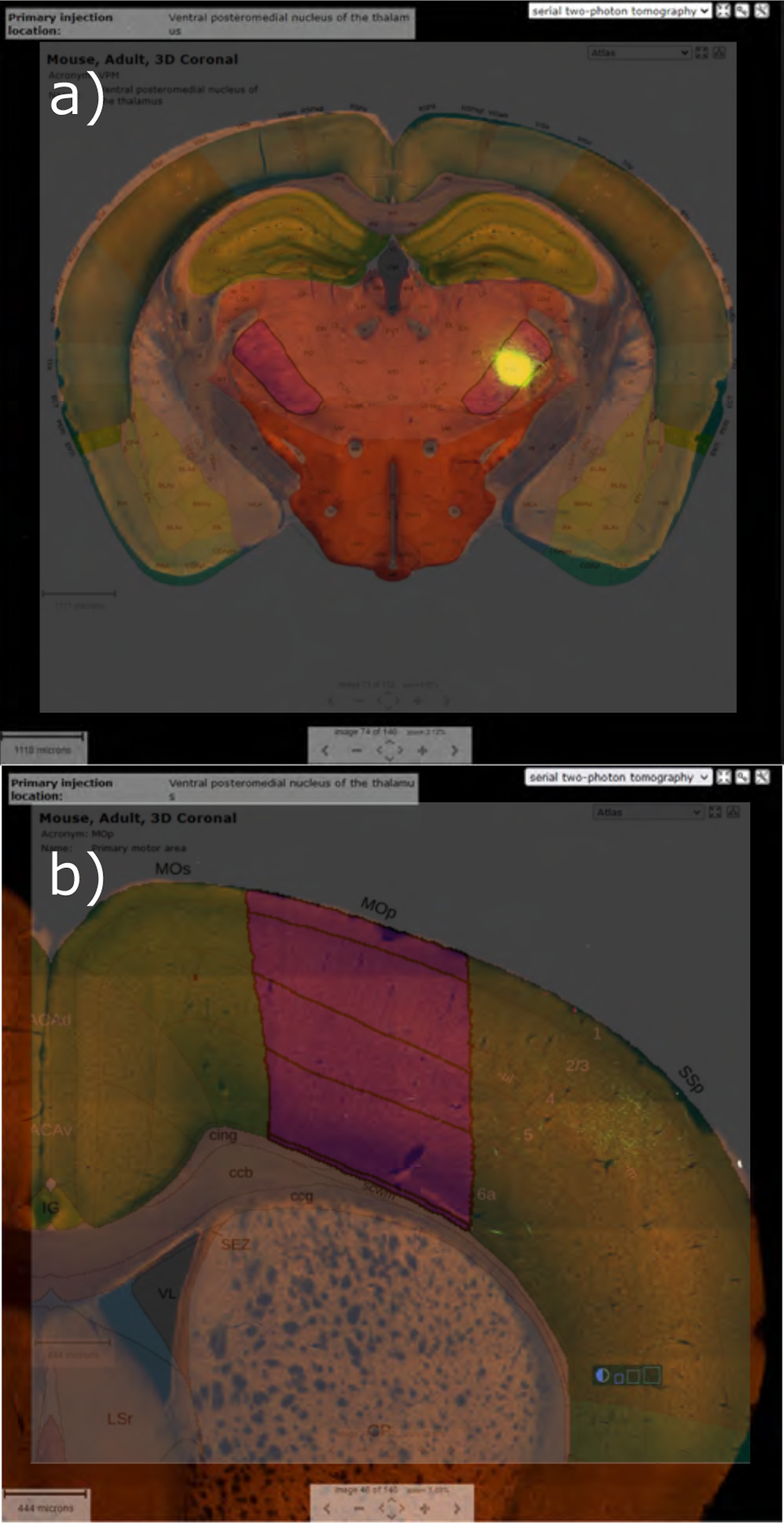
**a)** Injection structure VPM to **b)** MOp. Experiment number 312240825, Slc17a6-IRES-Cre. The shortcomings of experimental procedures are addressed in the discussion.

**Supplemental Fig. 34.**
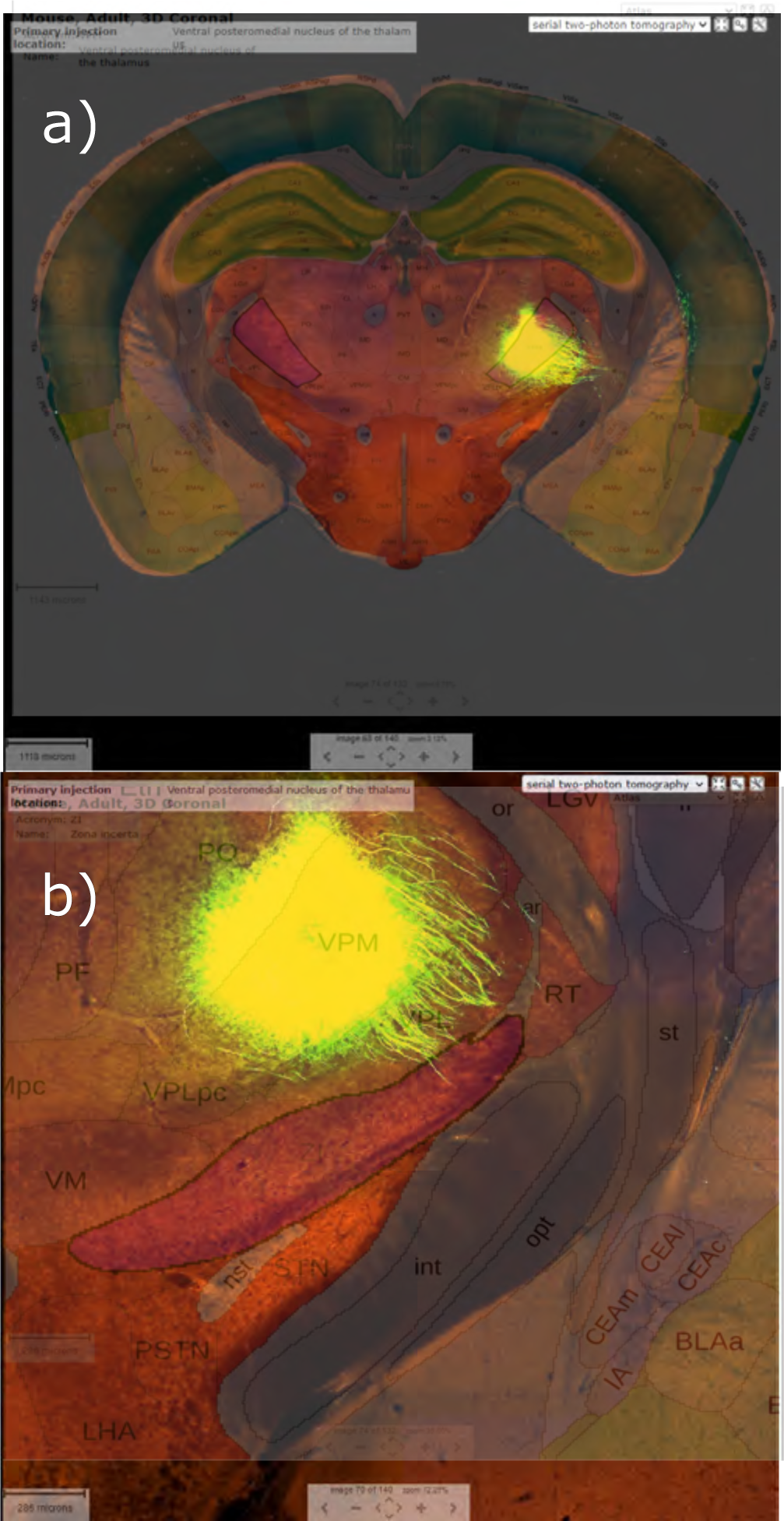
**a)** Injection structure VPM to **b)** ZI. Experiment number 268399868, Ppp1r17-Cre-NL146. The shortcomings of experimental procedures are addressed in the discussion.

**Supplemental Fig. 35.**
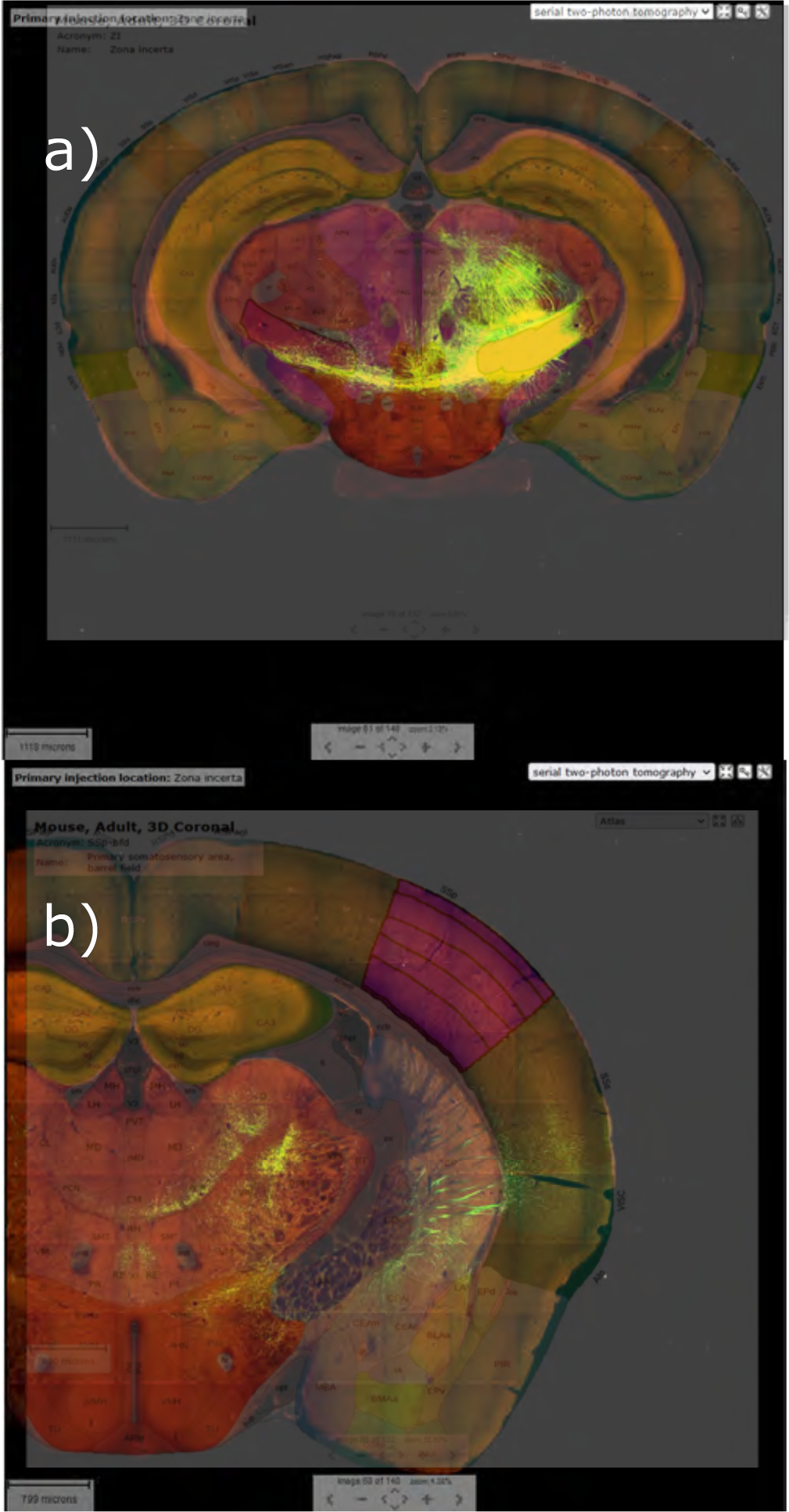
**a)** Injection structure ZI to **b)** bfd. Experiment number 113095845, wild-type. The shortcomings of experimental procedures are addressed in the discussion.

